# Stress-coping behavior during predator odor exposure is associated with differences in decision making

**DOI:** 10.64898/2026.05.05.722219

**Authors:** Brooke N Bender, Madison E Hoffman, Caroline G Krieman, Heavyn Smith, Joyce Besheer

## Abstract

Post-traumatic stress disorder (PTSD) and alcohol use disorder (AUD) are chronic psychiatric disorders that have overlapping symptomology and risk factors, including altered motivation and impulsive behavior. Inescapable exposure to a predator odor stressor (2,3,5-Trimethyl-3-Thiazoline (TMT)) produces PTSD-like symptomology in rats. Individual differences in stress-coping behaviors such as freezing and defensive digging during TMT exposure can predict long-term differences in alcohol-related behaviors and altered neurobiology. Here, we sought to evaluate the relationship between stress coping behavior during TMT exposure and different aspects of decision making. In Experiment 1, male and female rats were trained on an adjusting-amounts delay discounting task, and delay discounting curves were established before and >2 weeks after TMT exposure. In Experiment 2, female rats were trained to self-administer alcohol and sucrose in a concurrent choice procedure. Lever responses and preference for alcohol over sucrose were evaluated before and >2 weeks after TMT exposure, and then motivation for competing reinforcers was evaluated using progressive ratios. Active coping (digging) during TMT exposure was correlated with increased post-TMT impulsive choice (Experiment 1), reduced sucrose lever responses both before and after TMT exposure (Experiment 2), and reduced sucrose lever breakpoint (Experiment 2). Additionally, TMT-exposed rats had increased motivation for both alcohol and sucrose self-administration when available concurrently (Experiment 2). Overall, these findings suggest that behavior prior to and during a stressful experience can predict susceptibility to negative effects on decision making, which may help future studies identify the neurobiology underlying risk for aberrant reward-related behaviors after a traumatic event.

## 1. Introduction

Post-traumatic stress disorder (PTSD) and alcohol use disorder (AUD) are highly comorbid [1, 2]. Many factors may contribute to comorbidity between substance use disorders (SUDs) and PTSD, including increased risk of trauma in individuals with SUDs and increased risk of using substances to self-medicate trauma-induced negative affect [3, 4]. Additionally, overlapping genetic, behavioral, and biological factors may contribute to increased risk of developing both psychiatric disorders [1, 3].

Animal models of PTSD can be used to isolate and examine factors that contribute to comorbidity with AUD. Predator odor exposure is an ethologically relevant stressor that produces long-lasting behavioral and neurobiological effects that overlap with PTSD-related symptomologies [5]. Predator odor exposure usually involves a single exposure to a predator scent, such as predator urine, predator hair, or 2,3,5-Trimethyl-3-Thiazoline (TMT), a component of fox feces [5, 6, 7, 8]. Following predator odor exposure, studies have found increased home-cage alcohol drinking, alcohol self-administration, and sensitivity to alcohol in rodents [9, 10, 11, 12, 13, 14, 15]. Additionally, alcohol exposure can influence behavioral responses to predator odor exposure [12, 16]. Therefore, this model may be particularly useful for studying the overlap between PTSD and SUDs.

Additionally, animals exhibit diverse stress-coping responses to predator odor exposure, and several studies have shown that individual differences in these behavioral responses predict future drug-related behavior [16, 17, 18, 19, 20, 21]. Defensive digging (burying) and immobility (freezing) are common stress-coping responses displayed by rodents, including during exposure to predator odors [19, 22, 23]. We have previously shown that female rats that engaged in active-coping behaviors (more digging than immobility) self-administered more alcohol weeks after TMT exposure, whereas female and male rats that engaged in more passive-coping behaviors (more immobility than digging) did not [18]. This study revealed a relationship between digging and alcohol drinking, but whether or not pre-existing differences contributed to both behaviors remains unclear.

Several aspects of decision making that may contribute to drug use are impaired in both PTSD and AUD, including impulsivity and reward processing [24, 25]. Recent studies have indicated that impulsivity mediates the association between PTSD symptoms and scores on the Alcohol Use Disorders Identification Test (AUDIT) in trauma-exposed individuals, and impulsivity mediates the effects of childhood adversity and neglect on alcohol, nicotine, and cannabis use in young adulthood [26, 27]. Impulsive choice, one aspect of impulsivity, can be measured in both humans and rodents using delay discounting tasks, which evaluate how steeply the subjective value of a reward is discounted as the delay increases [28, 29]. We have shown that TMT exposure alters gene expression in several brain regions important for impulsive choice, including the nucleus accumbens and medial prefrontal cortex [29, 30, 31]. However, to date, the effects of predator odor exposure on delay discounting have not been studied.

Reward processing impairments in PTSD and AUD can influence choice between alcohol and non-drug rewards [32]. Heavy drinkers choose alcohol over food reward more than light drinkers [33], and greater AUD severity is associated with increased willingness to exert effort for alcohol over monetary reward [34]. Individuals with PTSD show reduced experience of pleasure derived from reward, or anhedonia [32, 35, 36, 37]. Additionally, individuals with comorbid PTSD and alcohol dependence show enhanced alcohol craving in response to drug-associated cues [38]. Together, these findings suggest an imbalance in the valuation of alcohol compared to non-drug rewards. Alcohol self-administration can be used to evaluate voluntary alcohol consumption in rodents, and the choice between alcohol and non-drug rewards can be modeled in concurrent self-administration paradigms, when distinct behaviors (e.g., pressing different levers) result in either alcohol or a non-drug reward, such as sucrose or a social interaction with a conspecific [39, 40, 41].

Here, we conducted two experiments to evaluate the relationship between stress-coping behavior during predator odor exposure and two different aspects of decision making that are impaired in AUD. In Experiment 1, rats were trained on a 2-lever delay-discounting task, and delay discounting for a sucrose reward was evaluated before and after TMT predator odor exposure. In Experiment 2, rats were trained on concurrent alcohol vs. sucrose operant self-administration, and self-administration and alcohol preference were evaluated before and after TMT exposure. Additionally, post-TMT motivation for alcohol and sucrose was measured in a progressive ratio test. In both experiments, stress-coping behaviors (digging and immobility) during TMT exposure were quantified, and the relationships between these behaviors and pre- and post-TMT decision making was evaluated. We hypothesized that TMT predator odor exposure would increase impulsive choice for a sucrose reward (Experiment 1) and choice of alcohol over sucrose (Experiment 2), and that these outcomes would be related to individual differences in stress-coping behaviors during TMT exposure.

## 2. Methods

### 2.1 Animals

Long-Evans rats arrived at 7 weeks of age. After 72 hours of undisturbed habituation, rats were handled for 1 minute daily for 5 days. Animals were housed in ventilated cages (Tecniplast, West Chester, PA) upon arrival to the facility and provided with *ad libitum* food and water unless otherwise specified. Rats were housed in temperature and humidity-controlled colony rooms with a 12-hour light-dark cycle. The facility lights turned on at 07:00 hours and off at 19:00 hours daily. All experiments were conducted during the light portion of the cycle except for overnight training. Animals were weighed daily (M-F). Rats were under continuous monitoring and care by the veterinary staff from the Division of Comparative Medicine at UNC-Chapel Hill. All experimental procedures were conducted in accordance with the NIH Guide for the Care and Use of Laboratory Animals and institutional guidelines.

### 2.2 Behavioral apparatus

Operant conditioning chambers (31 x 32 x 24 cm; Med Associates Inc., St. Albans, VT) were located in standard sound-attenuating cubicles equipped with an exhaust fan that provided ventilation and masked external noise. The operant conditioning chambers used in this study were previously described [18]. Briefly, all chambers had a single house light and tone generator. The left wall had 1 retractable lever that operated a syringe pump. The right wall had two retractable levers that operated a dipper arm. A white-cue light was located above each retractable lever.

### 2.3 TMT exposure

TMT exposure occurred in procedure rooms within the animal facility to allow for proper ventilation and air circulation. The materials and setup for the TMT exposures follow a previously described protocol [12, 13, 18, 19, 30, 42]. Chambers were equipped with ∼130g of ALPHA-dri^Ⓡ^ bedding evenly dispersed along the bottom of the chamber and a metal basket that hung from the right-side wall. Each metal basket contained a piece of filter paper onto which either 10μL of TMT (2,3,5-Trimethyl-3-Thiazoline, purity ≥ 97.0%, BioSRQ) or water were pipetted; the filter paper was inaccessible to the rat. Separate water (control) and TMT chambers and baskets were used so control animals were in an environment completely devoid of contact with TMT. Rats were separated into control and TMT groups based on pre-TMT impulsivity (Experiment 1) or alcohol vs sucrose preference (Experiment 2) such that the groups were balanced in their pre-TMT data. Rats were exposed up to 6 at a time, and only the rats to be exposed were present in the room. The TMT or water was pipetted onto the filter paper in each chamber, and the rat was placed inside shortly after. A video camera recorded the duration of the 10-minute exposure and all experimenters left the room for the session. Water control exposures (Experiment 1) occurred first, followed by TMT exposures. Due to changing lab protocols aimed at minimizing unnecessary stress to the control animals, in Experiment 2, the control animals were moved into the procedure room for 60 seconds. They remained in their home cages for this time, did not undergo water exposure, and were returned to the vivarium. Behavior recorded during the exposure was analyzed using ANY-maze Video Tracking Software [19]. Based on our previous work [12, 13, 18, 19, 30, 42], we focused on two primary behaviors during the TMT exposure: digging and immobility. Digging behavior was scored manually and analyzed in the software. The behavior was classified using the guidelines outlined in [23], and quantified in ANY-maze. Immobility behavior was operationally defined as a lack of movement for more than 2 seconds and this was assessed and quantified by the software. Total time spent digging and time spent immobile during the 10-minute exposure were the stress-coping measures used in analysis. After TMT exposure, rats were returned to the vivarium where they remained undisturbed (except for cage changes or to be weighed and fed when food restricted) for 2 weeks prior to returning to the assigned experiment.

### 2.4 Experiment 1

#### 2.4.1 Overview

In experiment 1, delay discounting for sucrose was evaluated both before and starting 2 weeks after TMT exposure. Male and female Long-Evans rats (n=24 per sex) were single-housed for the duration of the study and were food restricted to 85%-90% of their free-feeding body weight prior to the start of magazine training. One rat (male) was not food restricted for a portion of the study due to illness.

Operant conditioning chambers were programmed to deliver rewards via a dipper arm containing 0.1 ml of 10% (w/v) sucrose solution for 4 seconds. When multiple rewards were delivered, the dipper arm retracted for 1 second between 4-second deliveries. All lever responses and rewards were recorded, an infrared photobeam detected reward retrievals during the 4-second deliveries.

Training for this experiment was derived using methods and programs graciously provided from previous research that used delay-discounting in rats [43, 44].

#### 2.4.2 Magazine training

Subjects first completed magazine training, which occurred through ∼17-minute sessions daily for 4-6 days. During magazine training sessions, subjects received 30 sucrose solution deliveries via the dipper arm at an inter-trial interval of 15, 25, or 35-seconds until 30 trials were complete. Following each inter-trial interval, the house light turned on. After 10 seconds, the house light turned off and sucrose was delivered. All rats received at least 4 magazine training sessions.

#### 2.4.3 Overnight fixed ratio 1 (FR1) training

Upon completion of magazine training, rats underwent 2-5 15-hour overnight training sessions on an FR1 schedule, by which rats received a sucrose reward after one lever response. During overnight training, rats were free to respond on either the left or right active lever on the right wall of the self-administration chamber. Each trial began with a 10-second period when levers were retracted, the cue lights were off, and the chamber house light was on to signal trial start. After 10 seconds, both levers were inserted, cue lights above both levers turned on to signal sucrose availability, and the house light turned off. Upon a subject’s lever response, the opposite cue light was extinguished, and the dipper moved to the upward position for 4 seconds, allowing for sucrose retrieval. 10 seconds after the initial lever response, the following trial was initiated. Rats could receive up to 100 dippers per 15-hour overnight session. Rats that failed to retrieve >80% of rewards after the second overnight training were given additional magazine training sessions before proceeding to additional overnights.

#### 2.4.4 Trial training

Subjects underwent discrete trial training for 4-8 days. Similar to FR1 training, each trial began with the house light on to signal trial start, with levers retracted and cue lights off, for 10 seconds. The right and left levers were then inserted, and rats had 10 seconds to respond on either lever to initiate the choice phase of the trial. If subjects failed to respond on a lever to initiate the choice phase, a 10-second timeout with all lights off and levers retracted was initiated before the start of the next trial. Upon a lever response to initiate a choice trial, the house light turned off, and the cue lights turned on to signal sucrose availability. Subjects then had 10 seconds to choose to respond on either the left or right lever. Upon responding on either lever, the opposite cue light was extinguished and the dipper was activated for 4 seconds. Before the next trial began, rats underwent a timeout that lasted until 24 seconds after levers were inserted at trial start. Sessions ended after 30 total reinforcers were obtained or after 35 minutes elapsed.

Following trial training, subjects’ lever bias was analyzed to assign levers for subsequent delay-discounting sessions. The rat’s preferred lever on the final day of trial training was designated as the immediate lever, and the non-preferred lever was assigned as the delay lever.

#### 2.4.5 0-second delay training

Subjects underwent 0-second delay training for 9-18 days, or until the cohort’s average i value exceeded 4 for 2 of 3 consecutive days. During this training period, the reward delivered by the delay lever was not delayed. Similar to trial training, each trial began with the house light on to signal trial start, with levers retracted and cue lights off, for 10 seconds. The delay and immediate levers were then inserted, and rats had 10 seconds to respond on either lever to initiate the choice phase of the trial. If subjects failed to initiate the choice phase, a 10-second timeout with all lights off and levers retracted was initiated before the start of the next trial. Upon a lever response to initiate a choice trial, the house light turned off, and the cue lights turned on to signal sucrose availability. Subjects then had 10 seconds to choose between the delay and the immediate lever. Upon responding on either lever, the opposite cue light became extinguished, and the dipper was activated. If subjects responded on the delay lever, the dipper was activated 6 times. If subjects responded on the immediate lever, the dipper was raised 0-6 times. The dipper activation amount on the immediate lever (i value) was dependent on the subjects’ prior behavior. The value of the immediate lever started at 3, but increased by 1 after the delay lever was chosen and decreased by 1 after the immediate lever was chosen. This adjusting-amounts delay-discounting procedure, based on previous studies, allows for the determination of the indifference point, because subjects generally alternate between choices when the objective value of the immediate reward is relatively equal to the subjective value of the delayed reward within each subject [43, 44, 45, 46]. Before the next trial began, rats underwent a timeout that lasted until 50 seconds after levers were inserted at the start of the trial. If a subject made the same choice for 4 consecutive choice trials, the next choice trial was forced, in which the only lever inserted and the only cue light illuminated was the opposite of the previous 4 choices. Forced trials ensured subjects were attending to both levers and sampling both options, and these trials counted towards the total number of completed trials but did not influence i value. Sessions ended after 30 choice trials were completed or after 45 minutes had elapsed, whichever occurred first.

#### 2.4.6 Delay discounting

Following completion of the 0-second delay training, subjects began their pre-TMT delay discounting testing. These sessions were identical to 0-second delay training sessions, except that the delivery of the rewards on the delay lever was delayed by a fixed number of seconds within each session. Subjects completed a total of 4 sessions at each of the following delay intervals: 2 seconds, 4 seconds, 8 seconds, 16 seconds. When the delay lever was chosen, the dipper activated after the specified delay. Following TMT exposure and 2 weeks of undisturbed time, rats were re-tested on delay discounting as described for Pre-TMT Delay Discounting.

#### 2.4.7 Exclusions

One rat (female) was excluded due to death. One rat (female) was excluded because she failed to respond on the delay lever throughout 0-sec delay training. Eight rats (4 females, 4 males) were excluded after failing to retrieve more than 20 rewards during trial training session.

### 2.5 Experiment 2

#### 2.5.1 Overview

In experiment 1, choice self-administration between alcohol and sucrose was evaluated both before and starting 2 weeks after TMT exposure. Female Long-Evans rats (n=24) were utilized. Rats remained double-housed until the end of day 4 of sucrose self-administration training, after which they were moved to single housing in preparation for choice testing and TMT exposure.

Operant conditioning chambers were programmed such that a subject’s completion of the schedule on the left lever activated a syringe pump (Med Associates) that delivered 0.1 ml of solution into a liquid receptacle for 1.66 seconds. The syringe delivery was accompanied by illumination of a white cue light above the lever and a 1.66-s auditory tone. A subject’s completion of the schedule on the right lever activated a dipper arm that delivered 0.1 ml of solution for 4 seconds before retracting, which was accompanied by illumination of a white cue light above the lever.

#### 2.5.2 Alcohol self-administration training

Rats underwent two consecutive overnight alcohol self-administration training sessions. Rats were water-restricted for 24 hours leading up to the start of the first overnight. Following the first overnight session, rats were given 1 hour of water access, then resumed water restriction before the second overnight session. Rats had *ad libitum* access to water for the remainder of the study. During all alcohol self-administration sessions, only the retractable lever on the left wall that operated the syringe pump was inserted into the chamber. During the overnight sessions, 2% alcohol (volume/volume) and 10% sucrose (weight/volume) solution was delivered on an FR1 schedule until 50 reinforcers were obtained, and then the response requirement increased to FR2. Sessions ended after 16 hours or when rats earned a maximum of 480 reinforcers.

Following the two overnight sessions, rats were trained on alcohol self-administration on an FR2 schedule in 30-minute sessions using a sucrose-fading procedure in which alcohol concentration was gradually increased, and sucrose solution was gradually decreased until reaching a concentration of 15% alcohol, 0% sucrose. The exact order of sucrose fading is previously described (Ornelas et al., 2022). Rats remained at each concentration for 1 day, except for the 20% alcohol/0% sucrose phase, which was administered for 3 days. Following fading, the left lever administered 15% alcohol/0% sucrose for the remainder of the study. No inactive lever was inserted. Rats underwent 15% alcohol training for 10 days to gather baseline alcohol self-administration data.

#### 2.5.3 Sucrose self-administration training

Rats were then trained to self-administer sucrose, with only the rear lever on the right wall inserted (the alcohol lever on the left wall retracted). Rats underwent one sucrose overnight. The dipper arm delivered a 0.3% sucrose solution on an FR1 schedule until 50 reinforcers were obtained, and then the response requirement increased to FR2. The session terminated after 15 hours or upon reaching 200 sucrose reinforcers. Following the sucrose overnight, rats began sucrose training in 30-minute sessions, where the sucrose concentration was gradually increased until the collective average number of sucrose lever responses resembled the collective average of alcohol lever responses observed during the last three days of alcohol self-administration training. The exact order of sucrose concentration was 0.3% sucrose for 5 days, 0.5% sucrose for 3 days, and 1.0 % sucrose for 2 days before beginning alcohol versus sucrose choice training.

#### 2.5.4 Alcohol vs. sucrose choice self-administration

Upon completion of alcohol and sucrose self-administration training, rats began choice self-administration. During this phase, both the alcohol and sucrose levers were inserted, and the rats could choose which levers to respond on in a 30-minute session. Left lever responses delivered a 15% alcohol solution. Initially, right lever responses delivered 1.0 % sucrose solution. Sucrose concentration was maintained at 1.0% for 3 days, then increased to 1.5% for 1 day, and finally to 2.0% for the remainder of the pre-TMT portion of the study. The increase of sucrose was done to achieve similar average lever responses for sucrose and alcohol (average alcohol preference ∼0.5). Prior to TMT exposure, rats underwent choice self-administration for 9 days with the left lever at 15% alcohol and the right lever at 2% sucrose. Following a two-week break after TMT exposure, rats were returned to the choice procedure. Conditions remained the same as the pre-TMT choice, with rats beginning at 15% alcohol vs 2% sucrose for 10 days. Rats were then tested at 15% alcohol vs 1% sucrose for 12 days.

#### 2.5.5 Progressive-ratio (PR) choice testing

A final choice session was conducted using a progressive ratio 1 (PR1) schedule to measure subjects’ motivation for 15% alcohol versus 1% sucrose. Upon each completed response requirement on either the left or right lever, the response requirement for that lever increased by one (i.e., 1, 2, 3, etc.). Each lever operated on an independent PR1 schedule, such that responding on one lever did not affect the response requirement on the other. The progressive-ratio session lasted 30 minutes, and upon completion, the final ratio for each lever was recorded as the breakpoint, representing the maximum effort the subject was willing to expend to obtain the corresponding reinforcer.

### 2.6 Quantification and statistical analyses

#### 2.6.1 Calculation of i values and generation of delay discounting curves

The indifference point (i value) for each subject for each delay was calculated using a custom MATLAB (Math-works, Natick, MA, USA) script by averaging the value of the immediate lever in the last 10 choice trials on the final (4th) session at each delay. In 2 instances, the data from the 3rd session were used because the dipper arm was stuck up or the sucrose trough was absent from the chamber during the 4th session. Delay discounting curves were fit to i values using the Mazur hyperbolic model as previously described (*V* = *A*/(1+*kD*)) [43, 47]. *V* is the subjective value of the delayed reward (the i value), *A* is the initial (objective) value (6 sucrose rewards), *D* is the delay in seconds, and *k* is the calculated rate of discounting. Area under the curve was calculated using the i values at each delay.

#### 2.6.2 Alcohol preference

Alcohol preference was calculated by dividing the number of alcohol lever responses by the number of sucrose lever responses in each session.

#### 2.6.3 Median splits

The median digging value was used to separate animals based on individual differences in digging during TMT exposure, where subjects with digging values above the median were placed in the high-digging group, and subjects with digging values below the median were placed in the low-digging group. In Experiment 1, median splits were performed separately for each sex, so that there was the same number of males and females in low- and high-digging groups and any potential sex differences did not influence comparisons between groups.

#### 2.6.4 Statistical analyses

Behavioral data were recorded using MedPC software, and GraphPad Prism and SPSS were used to perform all statistical analyses. In Experiment 1, pre- and post-TMT delay discounting were analyzed with 3-way repeated-measures ANOVAs, with subjective value of the delayed reward (i value) as the dependent variable, sex and either TMT exposure group or digging group as between-subjects independent variables, and delay as a within-subjects independent variable. Delay discounting AUC and rate of discounting (k) were analyzed with 3-way repeated-measures ANOVAs, with AUC or rate of discounting (k) as the dependent variable, sex and either TMT exposure group or digging group as between-subjects independent variables, and pre-post TMT as a within-subjects independent variable. Delay discounting training data were analyzed with 2-way ANOVAs, with the training measure (such as reward retrievals, trials completed, etc.) as the dependent variable and sex and digging group as between-subjects independent variable.

In Experiment 2, choice self-administration behaviors were analyzed separately for each sucrose concentration with 2-way repeated-measures ANOVAs, with alcohol preference, alcohol lever responses, or sucrose lever responses as the dependent variable, TMT exposure group or digging group as a between-subjects independent variable, and session as a within-subjects independent variable. Pre-post TMT choice self-administration was analyzed using 2-way repeated-measures ANOVAs, with alcohol or sucrose lever responses as the dependent variable, TMT exposure group or digging group as a between-subject independent variable, and pre-post TMT as a within-subjects independent variable. Progressive ratio choice self-administration was analyzed using 2-way repeated-measures ANOVAs, with lever responses or breakpoint ratio as the dependent variable, TMT exposure group or digging group as a between-subjects independent variable, and reinforcer as a within-subjects independent variable. Single-lever self-administration training data were analyzed separately for alcohol and sucrose training, similar to choice sessions, except that sucrose concentrations were combined for analysis.

Stress-coping behaviors in Experiment 1 were analyzed with 3-way repeated-measures ANOVAs with either digging or immobility as the dependent variable, sex and TMT exposure group as between-subjects independent variables, and time as a within-subjects independent variable. Stress-coping behaviors in Experiment 2 were analyzed with 1-way repeated-measures ANOVAs with either digging or immobility as the dependent variable and time as a within-subjects independent variable. In both experiments, Pearson’s correlations were used to detect correlations between digging, immobility, and behavioral measures.

When there was no main effect of sex or an interaction with sex in Experiment 1, males and females were combined for final analysis with a 2-way repeated-measures ANOVA. When there was an interaction between variables, Tukey’s or Sidak’s post-hoc multiple comparisons analyses were performed when specified. For repeated-measures ANOVAs, Geisser-Greenhouse corrections were used to account for potential lack of sphericity. Significance was set at p<0.05 for all statistical analyses.

## 3. Results

### 3.1 Experiment 1: Impulsive choice for sucrose

Rats were trained on a delay discounting task (n=38; 18 females and 20 males included, see methods for exclusion criteria). During all phases of training, there were no differences between groups based on sex or future TMT exposure group (Supplementary Results; Supplementary Fig S1). After training, a pre-TMT delay discounting curve was established, then rats underwent TMT exposure or control. After a 2-week break, a post-TMT delay discounting curve was established (Fig 1A).

**Figure 1:**
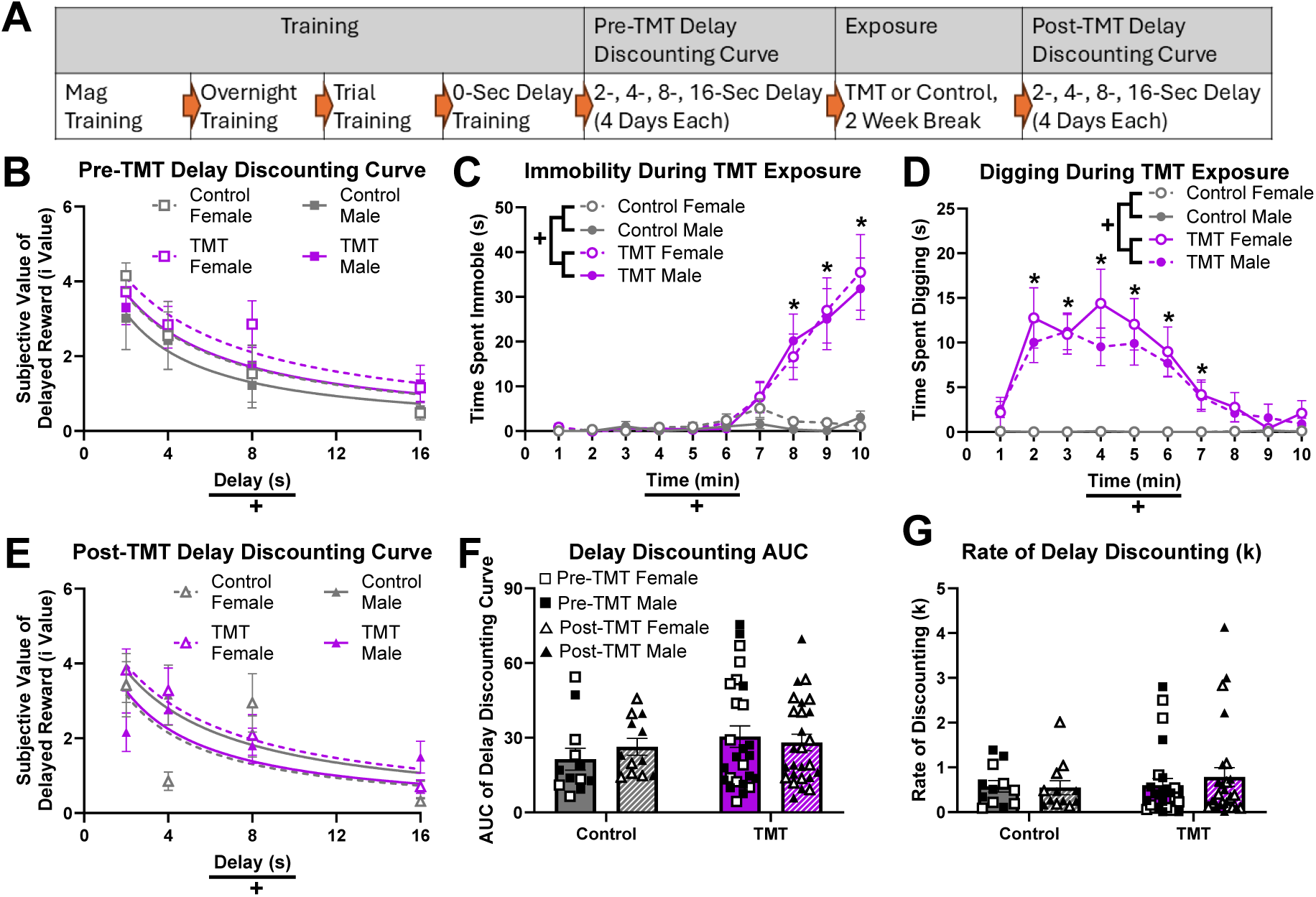
TMT exposure does not affect delay discounting in all TMT-exposed male and female rats Timeline of Experiment 1 (A). Males (closed symbols) and females (open symbols) are shown separately but were combined for analysis (B-D; F-G) unless there was a main effect of sex or interaction between sex and other variables (E). Main effects of delay (in seconds) on the subjective value of the delayed reward (i value) before (B) and after (E) TMT exposure are indicated by +. Main effects of time and TMT exposure on immobility (C) and digging (D) during TMT exposure are indicated by +. Post-hoc differences between TMT-exposed and control rats are indicated by * (C-D). Mazur’s hyperbolic equation was used to calculate a curve fit to delay discounting data (B, E). There were no main effects of sex, TMT group, or TMT exposure or interactions between variables for the AUC (area under the curve) of the delay discounting curve (F) or for the rate of delay discounting (k value) calculated based on Mazur’s hyperbolic equation (G). Graphs show group means ± SEM.

#### 3.1.1 TMT exposure does not impact impulsive choice in all TMT-exposed rats

Prior to TMT exposure, a delay discounting curve was established by determining the subjective value of the delayed reinforcer (i value) for delays of 2, 4, 8, and 16 seconds. There was no main effect of sex on i value (F_(1, 34)_=0.655, p = 0.424; 3-way rmANOVA), so males and females were combined for analysis. There was a main effect of delay on i value (F_(2.552, 91.87)_=21.28, p<0.001), where i value decreased as delay increased, but there was no main effect of future TMT exposure group (F_(1, 36)_=1.096, p = 0.302) or interaction between delay and future TMT exposure group (F_(2.552, 91.87)_=0.734, p = 0.514; 2-way rmANOVA; Fig 1B). These results suggest that delay discounting was expressed throughout all groups.

After baseline delay discounting was measured, rats underwent a 10-minute TMT exposure (n=26; 12 females and 14 males) or control (n=12; 6 females and 6 males). Digging and immobility behavior was quantified for each animal. There was no main effect of sex on immobility (F_(1, 34)_=0.059, p = 0.810) or digging (F_(1, 34)_=0.209, p = 0.651; 3-way rmANOVAs), so males and females were combined for analyses. For immobility during TMT exposure, there was a main effect of time (F_(1.644, 59.18)_=13.02, p<0.001) and TMT (F_(1, 36)_=12.90, p = 0.001), and there was an interaction between time and TMT (F_(1.644, 59.18)_=11.41, p < 0.001; 2-way rmANOVA; Fig 1C). Post-hoc analyses indicated that rats undergoing TMT exposure were immobile longer than control rats during minutes 8-10 (p<0.05; Sidak’s multiple comparisons test; Fig 1C). For digging during TMT exposure, there was a main effect of time (F_(3.204, 115.3)_=6.23, p < 0.001) and TMT (F_(1, 36)_=28.95, p < 0.001), and there was an interaction between time and TMT (F_(3.204, 115.3)_=6.334, p < 0.001; 2-way rmANOVA; Fig 1D). Post-hoc analyses indicated that rats undergoing TMT exposure were digging longer than control rats during minutes 2-7 (p < 0.05; Sidak’s multiple comparisons test; Fig 1D).

Two weeks after TMT exposure, a post-TMT delay discounting curve was established. There was a main effect of delay (F_(3, 102)_=16.52, p<0.001) on i value as well as an interaction between delay and sex (F_(3, 102)_=2.741, p = 0.047) and a 3-way interaction between delay, sex, and TMT exposure group (F_(3, 102)_=2.804, p = 0.044; 3-way rmANOVA; Fig 1E). There were no main effects of sex (F_(1, 34)_=0.002, p = 0.966) or TMT exposure group (F_(1, 34)_=0.180, p = 0.674), and there were no interactions between delay and TMT exposure group (F_(3, 102)_=2.444, p = 0.068) or sex and TMT exposure group (F_(1, 34)_=1.359, p = 0.252). Despite the 3-way interaction, there were no post-hoc differences in i value between male controls, female controls, male TMT, or female TMT rats at any of the 4 delays (Tukey’s multiple comparisons test).

To evaluate if there was a change in impulsive choice after TMT exposure compared to baseline, we calculated the area under the curve (AUC) and the rate of delay discounting (k value) for each rat and compared these values before and after TMT exposure. There was no main effect of sex on AUC (F_(1, 34)_=0.178, p = 0.676) or rate of delay discounting (k value) (F_(1, 34)_=0.0806, p = 0.778), so males and females were combined for analyses. For AUC, there were no main effects of TMT exposure group (F_(1, 36)_=1.159, p = 0.289) or pre-post TMT (F_(1, 36)_=0.1206, p = 0.730), and there was no interaction between TMT exposure group and pre-post TMT (F_(1, 36)_=0.977, p = 0.330; 2-way rmANOVA; Fig 1F). For rate of delay discounting (k value), there were no main effects of TMT exposure group (F_(1, 36)_=0.365, p = 0.550) or pre-post TMT (F_(1, 36)_=0.164, p = 0.688), and there was no interaction between TMT exposure group and pre-post TMT (F_(1, 36)_=0.317, p = 0.577; 2-way rmANOVA; Fig 1G). Overall, these results suggest that there were no differences in impulsive choice before or after TMT exposure when controls were compared to all TMT-exposed rats.

#### 3.1.2 Digging during TMT exposure correlates with post-TMT measures of impulsive choice

To determine if there were any correlations between digging and immobility behavior and delay discounting, we performed Pearson correlations between these behaviors and the pre-TMT i value on the final day of no delay, the pre- and post-TMT i values on the final day of each delay, AUC, and discounting rate (k value) (Fig 2A; Supplementary Table 1). There were negative correlations between time spent digging during TMT and post-TMT 4-sec i value (r(24)=-0.527, p = 0.006; Fig 2B), post-TMT 8-sec i value (r(24)=-0.436, p = 0.026; Fig 2C), and post-TMT AUC (r(24)=-0.469, p = 0.016; Fig 2D). There was also a positive correlation between time spent digging during TMT exposure and post-TMT discounting rate (k value) (r(24)=0.390, p = 0.049; Fig 2E). These results suggest that increased digging during TMT exposure was correlated with greater impulsive choice after TMT exposure. There were no correlations between digging in control rats with any of these measures.

**Figure 2:**
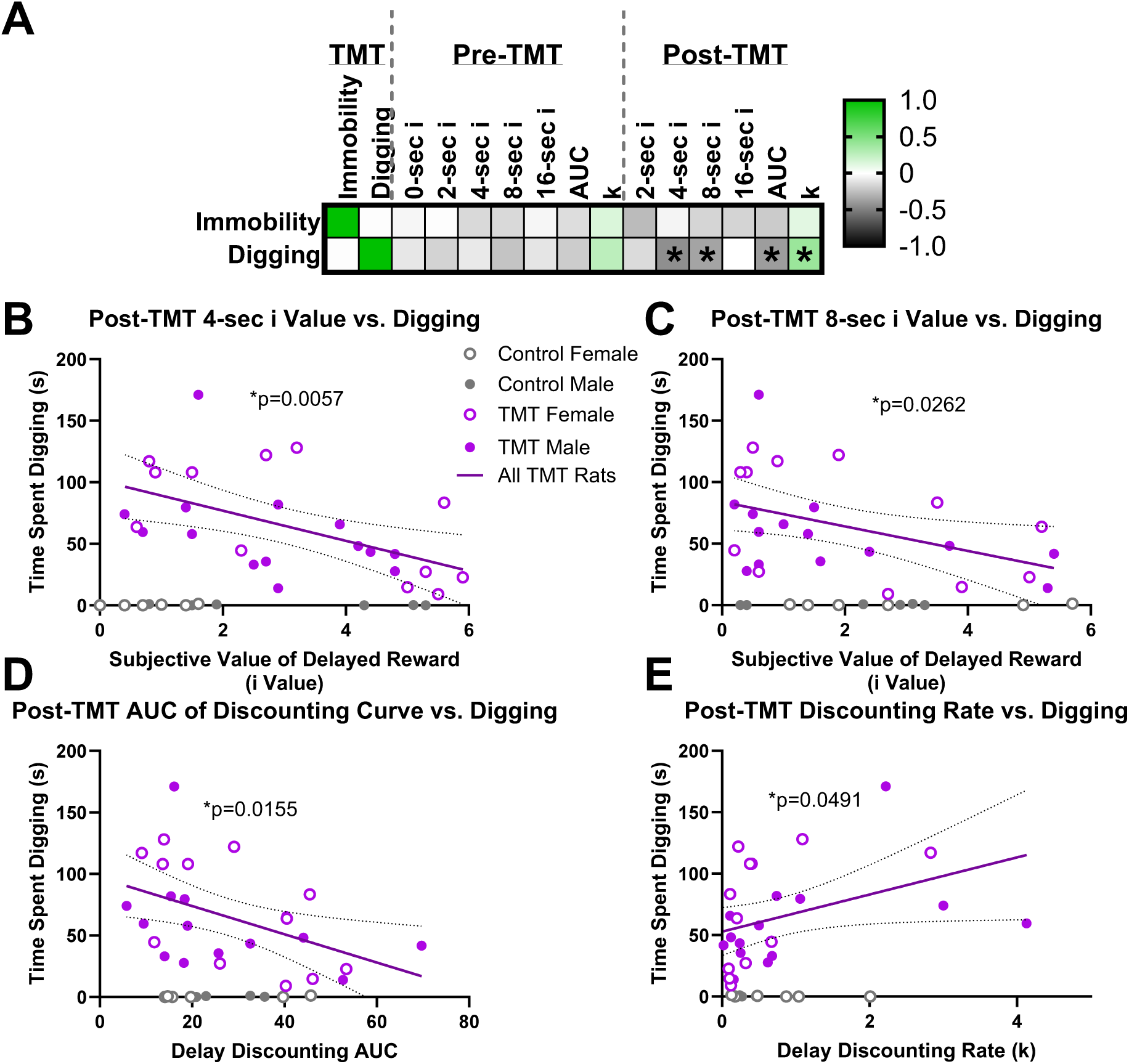
Digging during TMT exposure correlates with delay discounting after TMT exposure A Pearson correlation matrix (A) shows correlations between immobility and digging behavior during TMT exposure with various measures of delay discounting before and after TMT exposure, and significant correlations are indicated by * (A). For TMT-exposed rats but not control rats, there were negative correlations between digging during TMT and post-TMT 4-second i value (B), post-TMT 8-second i value (C), and Post-TMT AUC of the delay discounting curve (D). There was also a positive correlation between digging and the discounting rate (k value) for TMT-exposed rats but not control rats (E). Females (open symbols) and males (closed symbols) were combined for analysis. Graphs show lines of best fit with 95% confidence bands in dotted lines.

#### 3.1.3 Rats that dig more during TMT exposure (high digging) display more impulsive choice than rats that dig less (low digging)

Because digging during TMT exposure was correlated with post-TMT measures of delay discounting, we further examined the relationship between individual differences in digging and delay discounting. We separated TMT-exposed rats into low-digging and high-digging groups based on a median split within each sex of total time spent digging. For the pre-TMT i values, there was no main effect of sex (F(_1, 32_)=0.728, p = 0.400; 3-way rmANOVA), so males and females were combined. There was a main effect of delay on i value (F_(2.538, 88.84)_ = 21.65, p<0.001), but there was no main effect of digging group (F_(2, 35)_=1.278, p = 0.29) or interaction between delay and digging group (F_(5.076, 88.84)_=0.502, p = 0.777; 2-way rmANOVA; Fig 3A). For post-TMT i values, there was a main effect of delay (F_(2.732, 87.421)_=17.594, p < 0.001) and a main effect of digging group (F_(2, 32)_=3.835, p = 0.032), but there was no main effect of sex (F_(1, 32)_= 0.161, p = 0.691; 3-way rmANOVA; Fig 3B). There were also significant interactions between delay and digging group (F_(5.464, 32)=_2.596) and delay and sex (F_(2.732, 32)_=2.983), but there was no interaction between digging group and sex (F_(2, 32)_=0.862, p = 0.432) and there was no 3-way interaction (F_(5.464, 32)_=1.436, p = 0.215). Post-hoc analyses indicated that there were no differences between males and females at any delay, but at the 4-sec delay, low-digging rats had higher i values than both controls (p = 0.011) and high-digging rats (p = 0.016), and at the 8-sec delay, high-digging rats had lower i values than low-digging rats (p=0.025; Sidak’s multiple comparisons test; Fig 3B). These results suggest that although there were no differences before TMT exposure, delay discounting at intermediate delays (4 and 8 seconds) differed between low- and high-digging rats after TMT exposure.

**Figure 3:**
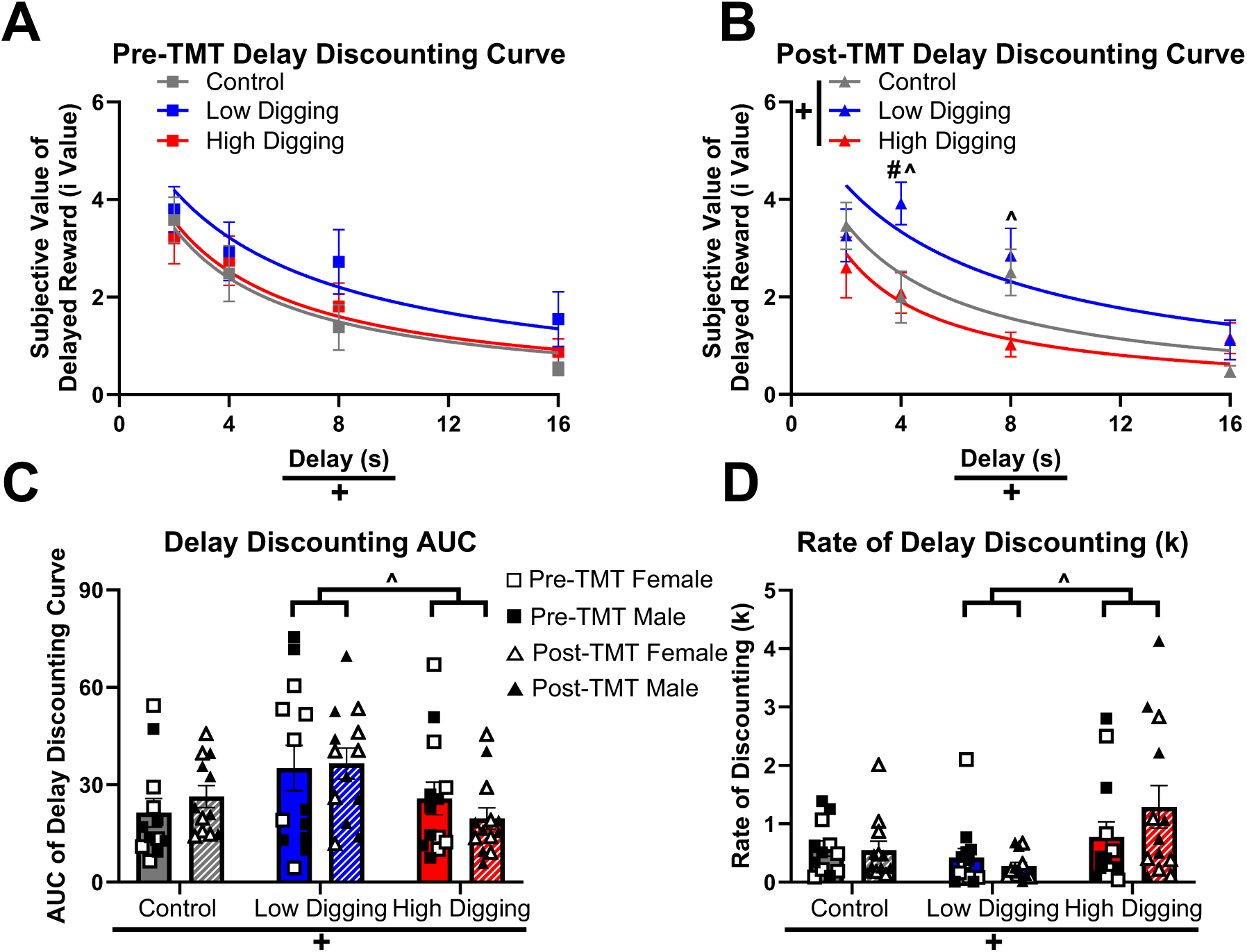
Individual differences in digging during TMT exposure reveal pre-existing differences in delay discounting that are exacerbated after TMT exposure To further investigate the relationship between digging during TMT exposure and delay discounting, we used a median split to separate TMT-exposed rats into low-digging and high-digging groups. Graphs collapse across sex for simplicity (A-B), but sex was considered as a variable. Males and females were combined for analysis if there was no main effect of sex or interaction between sex and other variables (A, C-D). Main effects of delay are indicated by + before (A) and after (B) TMT exposure. A main effect of digging group on the subjective value of the delayed reward (i value) after TMT exposure is indicated by + (B). Post-hoc differences between digging groups at each delay after TMT exposure are indicated as follows: # low digging vs. control and ^ low digging vs. high digging (B). Mazur’s hyperbolic equation was used to calculate a curve fit to delay discounting data (A, B). Main effects of digging group on AUC of the delay discounting curve (C) and the discounting rate (k value) (D) are indicated by +. Post-hoc differences between low-digging and high-digging groups on AUC of the delay discounting curve (C) and the discounting rate (k value) (D) are indicated by ^, but neither were different from controls. Graphs show group means ± SEM.

When comparing delay discounting after TMT exposure to baseline between controls, low-digging, and high-digging rats, there were no main effects of sex on AUC (F_(1, 32)_=0.394, p = 0.535) or discounting rate (k) (F_(1, 32)_=0.440, p = 0.512), so males and females were combined for analysis (3-way rmANOVAs). There was a main effect of digging group on AUC (F_(2, 32)_=3.503, p = 0.042), but there was no main effect of pre-post TMT (F_(1, 32)_=0.0002616, p = 0.9872) or interaction between digging group and pre-post TMT (F_(2, 32)_=0.836, p = 0.443; 2-way rmANOVA; Fig 3C). Post-hoc analyses indicated that high-digging rats had a lower AUC than low-digging rats (p = 0.045; Tukey’s multiple comparisons test; Fig 3C). There was a main effect of digging group on discounting rate (k) (F_(2, 32)_=4.883, p = 0.014), but there was no main effect of pre-post TMT (F_(1, 32)_=0.343, p = 0.562) or interaction between digging group and pre-post TMT (F_(2, 32)_=0.440, p = 0.512; 2-way rmANOVA; Fig 3D). Post-hoc analyses indicated that high-digging rats had a higher discounting rate (k) than low-digging rats (p = 0.009; Tukey’s multiple comparisons test; Fig 3D). Overall, these results suggest that although delay discounting in all TMT-exposed rats did not differ from controls before or after TMT exposure, high-digging rats displayed greater delay discounting than low-digging rats regardless of whether delay discounting was measured before or after TMT exposure, and this difference was particularly apparent at intermediate (4- and 8-sec) delays after TMT exposure. Additionally, results were similar between males and females.

### 3.2 Experiment 2: Choice between alcohol and sucrose self-administration

Rats (n=24; all female) were trained on a choice self-administration procedure where one lever was reinforced with alcohol and another was reinforced with sucrose solution. During all phases of training, there were no differences between groups based on future TMT exposure group or future digging group (Supplementary Results; Supplementary Fig S2). After initial choice self-administration training, rats underwent TMT exposure or control, and after a 2-week break, post-TMT choice self-administration was evaluated in standard choice sessions and in a choice session using progressive ratio schedules (Fig 4A).

**Figure 4:**
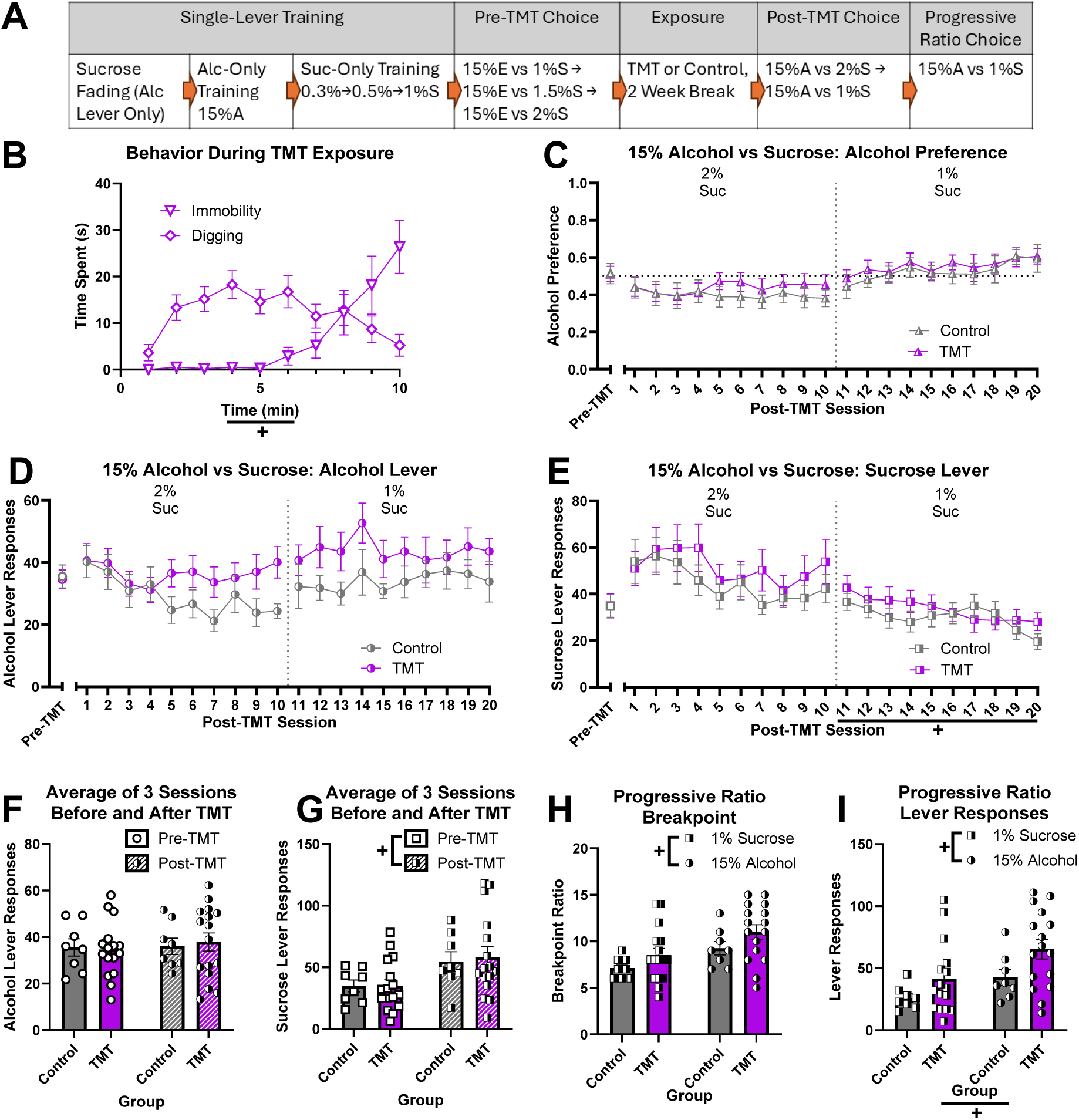
TMT exposure does not affect choice between self-administration of alcohol and sucrose but does increase responding on a progressive ratio schedule across reinforcers Timeline of Experiment 2 (A). Main effects of time on immobility and digging (B) in TMT-exposed rats during TMT exposure are indicated by +. Post-TMT alcohol preference (C), alcohol lever responses (D), and sucrose lever responses (E) during alcohol vs. sucrose choice sessions are shown with the average of the last 3 pre-TMT days to the left of an x axis break. A main effect of session on sucrose lever presses for 15% alcohol vs. 1% sucrose choice sessions (post-TMT) is indicated by + (E). The average number of alcohol lever responses (F) and sucrose lever responses (G) in the 3 sessions before (pre-TMT) and after (post-TMT) TMT exposure were compared, and a main effect of pre-vs-post TMT exposure on sucrose lever responses is indicated by + (G). Main effects of reinforcer (H, I) and TMT exposure (I) are indicated by + for breakpoint ratio (H) and lever responses (I) during a choice session where each reinforcer was delivered on its own individual progressive ratio schedule. Gray vertical dotted lines indicate a change in sucrose concentration (C-E), and a black horizonal dotted line at 0.5 alcohol preference indicates equal choice between alcohol and sucrose (C). Pre-TMT values are represented by open symbols, and post-TMT values are represented by half-shaded symbols. A or Alc = Alcohol; S or Suc = Sucrose. Graphs show group means ± SEM.

#### 3.2.1 TMT exposure does not affect choice between self-administration of alcohol and sucrose in females

After initial alcohol and sucrose choice self-administration training, rats underwent a 10-minute TMT exposure (n=16) or were left in their home cage as a control (n=8). During TMT exposure, there was a main effect of time on both immobility (F_(1.991, 29.87)_=10.49, p < 0.001) and digging (F_(1.039, 60.58)_=5.976, p < 0.001; 1-way rmANOVAs; Fig 4B). Post-TMT alcohol preference, alcohol lever responses, and sucrose lever responses were compared between controls and all TMT-exposed rats across 10 choice sessions between 15% alcohol and 2% sucrose and another 10 sessions when the sucrose concentration was reduced to 1%. For post-TMT 15% alcohol vs. 2% sucrose sessions, there were no main effects of session (F_(1.542, 33.92_=0.450, p = 0.591) or TMT exposure group (F_(1, 22)_=0.290, p = 0.595) on alcohol preference, and there was no interaction (F_(1.542, 33.92)_=0.693, p = 0.471; 2-way rmANOVA; Fig 4C). Additionally, for post-TMT 15% alcohol vs. 1% sucrose sessions, there were no main effects of session (F_(1.743, 38.34)_=3.223, p = 0.057) or TMT exposure group (F_(1, 22)_=0.152, p = 0.700) on alcohol preference, and there was no interaction (F_(1.743, 38.34)_=0.216, p = 0.777; 2-way rmANOVA; Fig 4C). For post-TMT 15% alcohol vs. 2% sucrose sessions, there were no main effects of session (F_(2.583, 56.83)_=1.98, p = 0.135) or TMT exposure group (F_(1, 22)_=1.621, p = 0.216) on alcohol lever responses, and there was no interaction (F_(2.583, 56.83)_=1.290, p = 0.286; 2-way rmANOVA; Fig 4D). Additionally, for post-TMT 15% alcohol vs. 1% sucrose sessions, there were no main effects of session (F_(2.303, 50.66)_=0.662, p = 0.541) or TMT exposure group (F_(1, 22)_=1.969, p = 0.175) on alcohol lever responses, and there was no interaction (F_(2.303, 50.66)_=0.349, p = 0.737; 2-way rmANOVA; Fig 4D). For post-TMT 15% alcohol vs. 2% sucrose sessions, there were no main effects of session (F_(1.557, 34.25)_=3.237, p = 0.063) or TMT exposure group (F_(1, 22)_=0.351, p = 0.560) on sucrose lever responses, and there was no interaction (F_(1.557, 34.25)_=0.657, p = 0.488; 2-way rmANOVA; Fig 4E). Additionally, for post-TMT 15% alcohol vs. 1% sucrose sessions, there was a main effect of session on sucrose lever responses (F_(1.422, 31.29)_=5.964, p = 0.012), but there was no main effect of TMT exposure group (F_(1, 22)_=0.262, p = 0.614) or interaction (F_(1.422, 31.29)_=1.906, p = 0.174; 2-way rmANOVA; Fig 4E).

To determine if alcohol vs. sucrose choice self-administration was altered after TMT exposure compared to baseline, we compared the average number of alcohol and sucrose lever responses in the 3 choice sessions prior to TMT exposure to the first 3 choice sessions that occurred 2 weeks after TMT exposure. There were no effects of TMT exposure group (F_(1, 22)_=0.0074, p = 0.932) or pre-post TMT (F_(1, 22)_=0.939, p = 0.343) on alcohol lever responses, and there was no interaction (F_(1, 22)_=498, p = 0.488; 2-way rmANOVA; Fig 4F). For sucrose lever responses, there was a main effect of pre-post TMT (F_(1, 22)_=21.83, p < 0.001), but there was no main effect of TMT exposure group (F_(1, 22)_=0.033, p = 0.857) or interaction (F_(1, 22)_=0.1528, p = 0.700; 2-way rmANOVA; Fig 4G), where sucrose lever responses increased post-TMT (after the two-week break) regardless of whether rats were exposed to TMT. Together, these results suggest that TMT exposure did not impact alcohol preference, alcohol self-administration, or sucrose self-administration when both reinforcers were available concurrently on an FR2 schedule.

#### 3.2.2 TMT exposure increases responding for sucrose and alcohol on a progressive ratio schedule in females

After post-TMT choice self-administration sessions, rats underwent a final self-administration choice session where the alcohol and sucrose levers were reinforced on progressive ratio schedules to evaluate motivation when more effort was required to obtain reinforcers. Breakpoint ratio (the ratio at which each animal did not obtain a reinforcer), as well as lever responses for each reinforcer were compared. For breakpoint ratio, there was a main effect of reinforcer (F_(1, 22)_=9.925, p = 0.005), where both groups had a higher breakpoint for 15% alcohol than for 1% sucrose, but there was no effect of TMT exposure group (F_(1, 22)_=3.081, p = 0.093) or interaction (F_(1,22)_=0.065, p = 0.801; 2-way rmANOVA; Fig 4H). For lever responses, there was a main effect of reinforcer (F_(1, 22)_=8.250, p = 0.009) and a main effect of TMT exposure group (F_(1, 22)_=4.626, p = 0.0427), where both groups made more responses for alcohol and TMT-exposed rats made more responses in general than controls, but there was no interaction (F_(1, 22)_=0.276, p = 0.605; 2-way rmANOVA; Fig 4I). Together, these results suggest that both TMT-exposed and control rats were more motivated for 15% alcohol compared to 1% sucrose. Additionally, TMT-exposed rats were more motivated for both reinforcers compared to controls even over 6 weeks after TMT exposure.

#### 3.2.3 Behavior during TMT exposure correlates with pre- and post-TMT sucrose self-administration and post-TMT sucrose progressive ratio breakpoint in females

We performed Pearson correlations to determine if there were any correlations between digging and immobility behavior and alcohol vs. sucrose choice self-administration. We evaluated correlations between stress-coping behaviors and the average alcohol preference, alcohol lever responses, and sucrose lever responses during the 3 sessions pre-TMT and post-TMT. Additionally, we also evaluated correlations between stress-coping behavior and post-TMT lever responses and breakpoint ratios for sucrose and alcohol under a progressive ratio schedule of reinforcement (Fig 5A; Supplementary Table 2). There was a negative correlation between time spent digging during TMT exposure and both pre-TMT sucrose lever responses (r(14)=-0.502, p = 0.047; Fig 5B) and post-TMT sucrose lever responses (r(14)=-0.572, p = 0.021; Fig 5C). There was also a positive correlation between time spent immobile during TMT exposure and post-TMT sucrose lever responses (r(14)=0.549, p = 0.028; Fig 5D). Finally, there was a negative correlation between time spent digging during TMT exposure and breakpoint ratio for sucrose (r(14)=-0.526, p = 0.037; Fig 5E), which was only measured post-TMT at the end of the experiment. Together, these results suggest that animals that made fewer responses for sucrose prior to TMT dug more during TMT exposure, and this effect was maintained after TMT exposure. Furthermore, animals that dug more also had lower breakpoint ratios for sucrose, which suggests lower motivation for sucrose.

**Figure 5:**
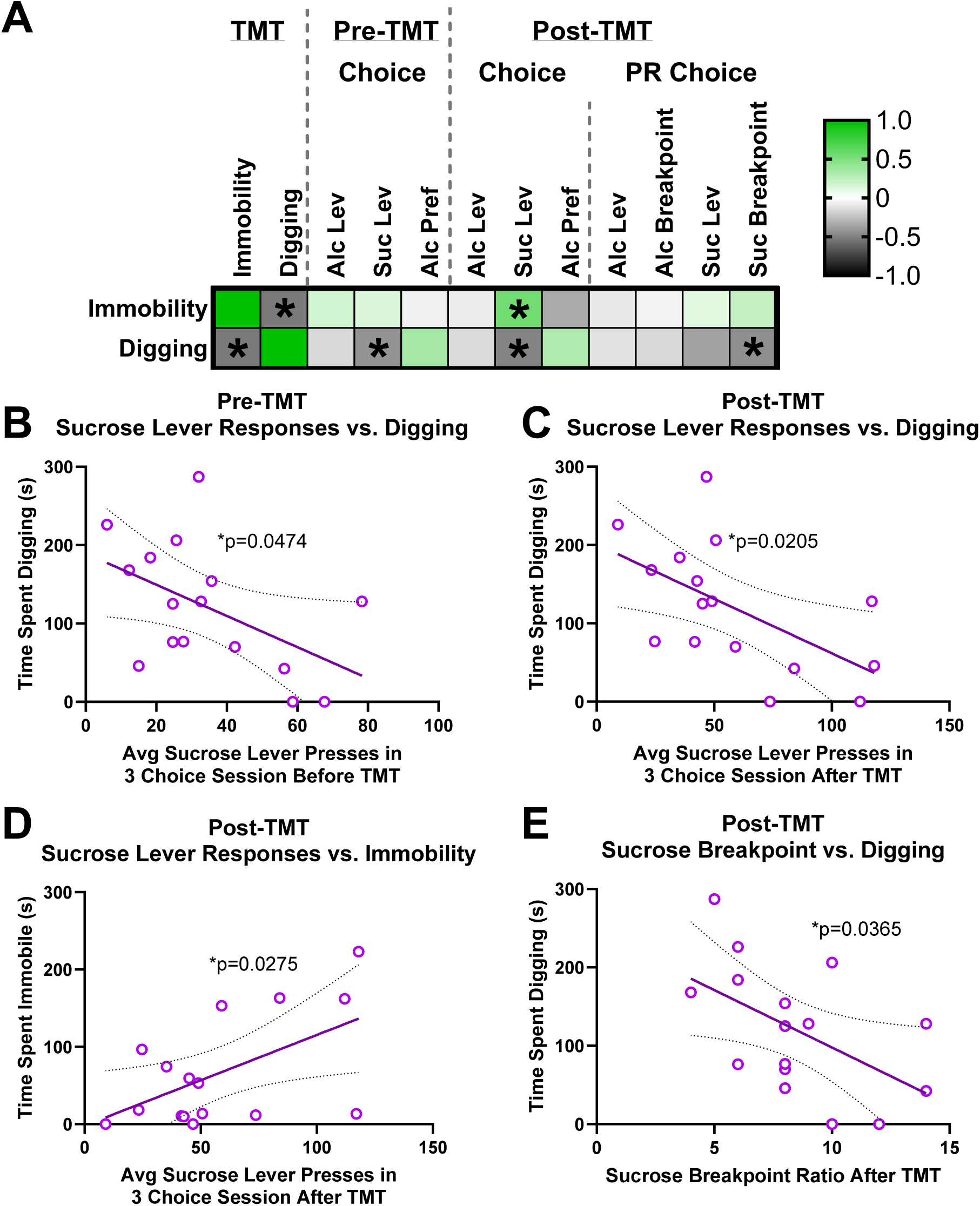
Behavior during TMT exposure correlates with pre- and post-TMT sucrose self-administration and sucrose breakpoint ratio A Pearson correlation matrix (A) shows correlations between immobility and digging behavior during TMT exposure with the average alcohol preference, alcohol lever responses, and sucrose lever responses during the 3 sessions before and after TMT exposure, as well as lever responses and breakpoint ratios for sucrose and alcohol under a progressive ratio schedule of reinforcement (post-TMT) (A). Significant correlations are indicated by * (A). There were negative correlations between digging during TMT exposure and pre-TMT sucrose lever responses (B), post-TMT sucrose lever responses (C), and post-TMT sucrose breakpoint ratio (E). There was also a positive correlation between immobility during TMT exposure and post-TMT sucrose lever responses (D). Graphs show lines of best fit with 95% confidence bands in dotted lines.

#### 3.2.4 There were no differences between low-digging and high-digging rats in choice of self-administration of alcohol and sucrose in females

Because digging during TMT exposure was correlated with responding for sucrose before and after TMT exposure, we used a median split of total time spent digging to separate animals into low-digging and high-digging groups (as in Experiment 1) to further investigate individual differences. There were no effects of session (F_(1.533, 32.19)_=0.730, p = 0.455) or digging group (F_(2, 21)_=0.203, p = 0.818) or interaction (F_(3.066, 32.19)_=0.588, p = 0.631) for post-TMT alcohol preference for 15% alcohol vs. 2% sucrose sessions, and there were also no effects of session (F_(1.713, 35.97)_=2.273, p = 0.056) or digging group (F_(2, 21)_=0.223, p = 0.802) or interaction (F_(3.426, 35.97)_=0.516, p = 0.698) for post-TMT alcohol preference for 15% alcohol vs. 1% sucrose sessions (2-way rmANOVAs; Fig 6A). There were no effects of session (F_(2.643, 55.49)_=1.684, p = 0.186) or digging group (F_(2, 21)_=1.888, p = 0.176) or interaction (F_(5.285, 55.49)_=1.202, p = 0.320) for post-TMT alcohol lever responses for 15% alcohol vs. 2% sucrose sessions, and there were also no effects of session (F_(2.479, 52.07)_=0.870, p = 0.445) or digging group (F_(2, 21)_=3.100, p = 0.066) or interaction (F_(4.959, 52.07)_=0.429, p = 0.825) for post-TMT alcohol lever responses for 15% alcohol vs. 1% sucrose sessions (2-way rmANOVAs; Fig 6B). For post-TMT sucrose lever responses, there was a main effect of session for 15% alcohol vs. 2% sucrose (F_(1.533, 32.20)_=3.732, p = 0.045) and 15% alcohol vs. 1% sucrose (F_(1.410, 29.60)_=6.963, p = 0.007), but there was no effect of digging group (F_(2, 21)_=1.148, p = 0.337) or interaction (F_(3.067, 32.20)_=1.259, p = 0.305) for 15% alcohol vs. 2% sucrose, and there was no effect of digging group (F_(2, 21)_=0.378, p = 0.690) or interaction (F_(2.819, 29.60)_=1.548, p = 0.224) for 15% alcohol vs. 1% sucrose (2-way rmANOVAs; Fig 6C). These results suggest that individual differences in digging behavior during TMT exposure were not related to post-TMT alcohol preference, alcohol self-administration, or sucrose self-administration across sessions.

**Figure 6:**
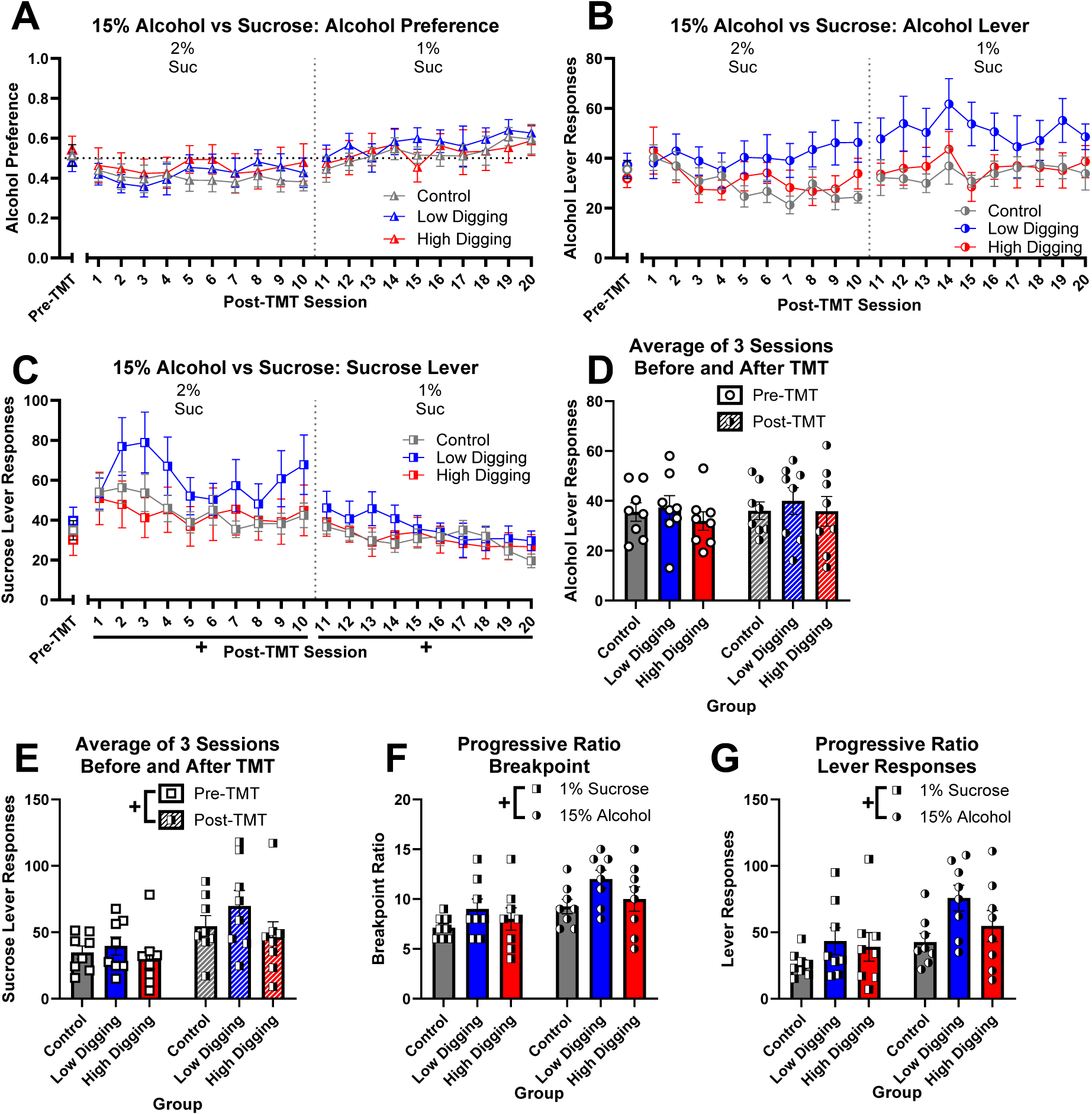
There were no differences between digging groups in alcohol or sucrose choice Post-TMT alcohol preference (A), alcohol lever responses (B), and sucrose lever responses (C) during alcohol vs. sucrose choice sessions are shown for controls, low-digging, and high-digging rats, with the average of the last 3 pre-TMT days to the left of an x axis break. Main effects of session on sucrose lever presses for 15% alcohol vs. 2% sucrose choice and 15% alcohol vs. 1% sucrose choice sessions (post-TMT) are indicated by + (C). The average number of alcohol lever responses (D) and sucrose lever responses (E) in the 3 sessions before (pre-TMT) and after (post-TMT) TMT exposure were compared between digging groups, and a main effect of pre-vs-post TMT exposure on sucrose lever responses is indicated by + (E). Main effects of reinforcer are indicated by + for breakpoint ratio (F) and lever responses (G) during a choice session using a progressive ratio schedule. Gray vertical dotted lines indicate a change in sucrose concentration (A-C), and a black horizonal dotted line at 0.5 alcohol preference indicates equal choice between alcohol and sucrose (A). Pre-TMT values are represented by open symbols, and post-TMT values are represented by half-shaded symbols. Graphs show group means ± SEM.

Additionally, to determine if alcohol vs. sucrose choice was altered differentially based on digging behavior after TMT exposure compared to baseline, we compared the average number of alcohol and sucrose lever responses in the 3 choice sessions prior to TMT exposure to the first 3 choice sessions that occurred 2 weeks after TMT exposure. For alcohol lever responses, there were no effects of digging group (F_(2, 21)_=0.313, p = 0.735) or pre-post TMT (F_(1, 21)_=1.563, p = 0.225), and there was no interaction (F_(2, 21)_=0.277, p = 0.761; 2-way rmANOVA; Fig 6D). For sucrose lever responses, there was a main effect of pre-post TMT (F_(1, 21)_=26.73, p < 0.001), but there was no effect of digging group (F_(2, 21)_=1.063, p = 0.363) or interaction (F_(2, 21)_=0.910, p = 0.418; 2-way rmANOVA; Fig 6E), where sucrose lever responses increased post-TMT (after the two-week break) regardless of TMT exposure or digging group.

During a final choice session when alcohol and sucrose were delivered on individual progressive ratio schedules, there was a main effect of reinforcer on breakpoint ratio (F_(1, 21)_=11.42, p = 0.003), where all groups had a higher breakpoint for 15% alcohol than 1% sucrose, but there was no effect of digging group (F_(2, 21)_=2.754, p = 0.087) or interaction (F_(2, 21)_=0.200, p = 0.820; 2-way rmANOVA; Fig 6F). Additionally, there was a main effect of reinforcer on lever responses (F_(1, 21)_=10.46, p = 0.004), where all groups made more lever responses for 15% alcohol than for 1% sucrose, but there was no effect of digging group (F_(2, 21)_=3.180, p = 0.062) or interaction (F_(2, 21)_=0.655, p = 0.530; 2-way rmANOVA; Fig 6G). Altogether, these results suggest that although digging during TMT exposure was correlated with pre- and post-TMT sucrose lever responding and post-TMT motivation for sucrose, individual differences in digging behavior during TMT exposure (low and high-digging groups) were not related to differences in alcohol vs. sucrose choice self-administration behavior.

## 4. Discussion

In two experiments, we evaluated the effects of TMT predator odor exposure on aspects of decision making relevant to alcohol use disorders: impulsive choice measured by delay discounting (Experiment 1) and choice between self-administration of sucrose and alcohol (Experiment 2). Additionally, we evaluated how these aspects of decision making were related to stress-coping behavior displayed during TMT exposure. Experiment 1 showed that although impulsive choice was not impacted in all TMT-exposed rats, more digging during TMT exposure was correlated with subsequently increased impulsive choice, and high-digging rats made more impulsive choices than low-digging rats after the stressor, particularly at moderate (4-8-second) delays. Additionally, high-digging rats displayed higher rates of delay discounting than low-digging rats both before and after TMT exposure. These results suggest that greater rates of delay discounting predict active stress-coping responses to predator odor exposure, and active stress-coping responses predicted greater susceptibility to stress-induced increases in impulsive choice. Experiment 2 showed that although TMT exposure did not impact alcohol or sucrose concurrent choice self-administration or preference for alcohol over sucrose on an FR2 schedule, it did lead to increased lever responding for both sucrose and alcohol when available on a progressive ratio schedule, indicating an overall increase in motivation for reward. Additionally, more digging during TMT exposure was correlated with fewer lever responses for sucrose both before and after exposure, as well as reduced breakpoint ratio for sucrose on a progressive ratio schedule. Both of these results suggest that lower motivation for sucrose predicts more active stress-coping responses to predator odor exposure. Importantly, post-TMT decision making in both experiments was measured between 2-6 weeks after the TMT exposure, which suggests that both the effects of the stressor and associations with stress-coping behavior were long-lasting.

### 4.1 Experiment 1: Impulsive choice for sucrose

Despite clinical literature supporting a relationship between increased delay discounting and a history of trauma or chronic stress [25, 48, 49, 50], few preclinical studies have evaluated the effects of stress on impulsive choice in rodents. Chronic corticosterone exposure in adolescent male rats led to increased impulsive choice in a delay discounting task in adulthood [51]. Chronic unpredictable stress increased impulsive choice in male GDNF (glial-derived neurotrophic factor)-deficient heterozygous mice, which have impaired dopamine regulation in the striatum, but not in wild-type controls [52]. Additionally, in male rats, intermittent social defeat stress increased impulsive choice particularly in subjects with lower impulsive choice at baseline, and these effects persisted up to 2 weeks after the final social defeat session [53]. Our results similarly suggest that stress increases impulsive choice in a subset of subjects. When individual differences in stress-coping behavior (particularly digging) were not considered, there was no overall effect of TMT exposure on impulsive choice. However, effects emerged when we considered individual differences in stress-coping behavior. Our findings differ slightly from those previous [53] in that the group that increased impulsive choice after stressor exposure exhibited higher impulsive choice at baseline, which could be due to differences in the type or timing of the stressor. Together, the current study supports previous literature indicating that individual differences in genetics and behavior may contribute to stressor susceptibility in a subset of animals, which highlights the importance of examining individual differences in the effects of stress, especially on impulsive choice.

We measured baseline impulsive choice prior to TMT exposure to determine if relationships between stress-coping behavior and decision making were pre-existing or induced by the stressor. Interestingly, digging during TMT exposure, which is considered an active-coping strategy, was correlated only with measures of delay discounting after TMT exposure, and high-digging rats only showed significantly reduced subjective value of the delayed reward compared to low-digging rats after TMT exposure. Therefore, the differences in delay discounting depending on stress-coping strategy were most pronounced after the stressor. However, when we compared AUC of the delay discounting curve and the delay discounting rate before and after TMT exposure, high-digging rats had a smaller AUC and greater rate of discounting compared to low-digging rats in general. Importantly, there were no differences between digging groups in any measures of task learning, so these differences are unlikely to be attributed to issues with learning or differences in motivation to perform the task. These results point to pre-existing differences in impulsive choice that are exacerbated by predator odor stress in rats that display more active stress-coping behavior.

Stress-coping strategies have been previously associated with differences in impulsive choice, particularly in the Roman high- and low-avoidance rat lines selectively bred for active avoidance behavior [54, 55]. Roman low-avoidance rats typically show more freezing (passive-coping) responses to stressors, whereas Roman high-avoidance rats typically show more flight/escape (active-coping) responses [55]. In a study evaluating only females, Roman high-avoidance rats showed increased impulsive choice compared to the low-avoidance line [54]. Our results demonstrating increased impulsive choice in high-digging rats extend the association between active-coping behaviors and impulsive choice to both male and female Long-Evans rats. Together, these findings point to biological differences that may underlie both active-coping responses to stress and impulsive choice, which remain to be investigated.

We conducted delay discounting in both male and female Long-Evans rats. Despite some interactions between sex and other variables in post-TMT delay discounting, these interactions were driven by minor differences in changes in subjective value of the delayed reward across delays. Further, there were no sex-related differences between groups at any single delay or for the AUC of the delay-discounting curve or rate of delay discounting. Effects of sex and hormones on delay discounting have been reported in both humans and rodents [56, 57, 58, 59, 60]. In the present study we did not examine phases of estrous, and we used multiple days of testing that likely occurred across varying phases of the cycle. Together, this may have contributed to the lack of sex differences. These factors could be examined in follow-up studies.

Although we are interested in the overlap between impulsive choice, stress responsivity, and alcohol-related behaviors, it is important to note that the current experiment measured impulsive choice for a sucrose reward in animals with no exposure to alcohol. There is ample evidence for a relationship between alcohol-related behaviors and impulsive choice for natural rewards, such as money for humans and sucrose for rodents [34, 44, 61, 62, 63], but impulsive choice for sucrose may not always generalize to impulsive choice for alcohol. Additionally, we previously showed that rats extensively exposed to alcohol during alcohol discrimination training engaged in more digging during TMT predator odor exposure compared to alcohol-naïve rats [12]. Therefore, exposure to alcohol could also impact stress responsivity and susceptibility to stress-induced increases in impulsive choice, which could be investigated in future experiments.

### 4.2 Experiment 2: Choice between alcohol and sucrose self-administration

In the present study, there was no effect of TMT exposure on alcohol or sucrose lever responses, or alcohol preference in female rats when alcohol and sucrose were concurrently available for self-administration, even when rats were separated based on individual differences in digging. This was somewhat surprising given our previous findings that TMT exposure increased sweetened alcohol self-administration in male Long-Evans rats [9] and unsweetened alcohol self-administration in female Long-Evans rats that displayed more digging (active coping) than immobility (passive coping) in response to TMT exposure [18]. Other studies have also shown increases in alcohol self-administration after predator odor stressor exposure in male Wistar rats [10], as well as short-term increases in alcohol intake after predator odor stress in male and female C57BL/6J mice [11]. In the previous studies, no non-alcohol reward was concurrently available, so the presence of a concurrently available reinforcer (sucrose) may have contributed to the lack of effect on alcohol self-administration in the present study.

In a test conducted several weeks after the initial TMT exposure, TMT exposure increased lever responding for both sucrose and alcohol when animals were responding on a progressive ratio schedule, a measure of motivation for reward [64]. These results show that TMT predator odor exposure increases motivation for both alcohol and a natural sucrose reward, and this effect is persistent for several weeks. Although the effects of predator odor exposure on the self-administration of several drugs have been studied, the effects on self-administration of sucrose and other rewards are unclear. Here, we provide evidence that increased motivation following TMT exposure may be generalized between sucrose and alcohol, and future studies should examine if there is a common underlying mechanism.

Despite a general increase in motivation for sucrose after TMT exposure, a subset of rats that displayed more active stress-coping behavior (digging) made fewer lever responses for sucrose on a progressive ratio schedule as well as fewer sucrose lever responses during standard choice self-administration before and after predator odor stress. Together, these results suggest that the pre-existing correlation between more digging and fewer sucrose lever responses persisted despite a general increase in motivation for sucrose following TMT exposure. Interestingly, post-TMT lever responses for sucrose were also positively correlated with immobility, which suggests an inverse relationship between digging and immobility on post-TMT sucrose self-administration. Reduced sucrose self-administration and motivation for sucrose could indicate anhedonia [65]. Therefore, high-digging rats may display anhedonia-like behavior even prior to stressor exposure that persists after TMT exposure.

Despite a relationship between digging and lever responses for sucrose, there was no correlation between digging and lever responses for alcohol or preference for alcohol over sucrose. Although differences in motivation for a natural reward, like sucrose, in the presence of a drug like alcohol may influence choice between the two, the present results are difficult to interpret in the context of alcohol-related behavior. Rats in the present study could freely self-administer both sucrose and alcohol, and self-administration of one did not preclude subsequent self-administration of the other. It is possible that a different session structure, with more limited or discrete trials, would identify additional effects.

One unexpected finding was an increase in sucrose, but not alcohol, self-administration in both the unstressed controls and TMT-exposed groups after the two-week break from self-administration following predator odor exposure. This increase in sucrose self-administration could be due to incubation of craving for sucrose, which has been demonstrated in rats 7 and 30 days after abstinence from sucrose [66]. Although incubation of craving in rats has also been shown for alcohol, one study indicated that responding on the alcohol lever was significantly elevated after 8, but not 4 weeks following abstinence [67]. Therefore, it is likely that the 2-week break in the present study was sufficient to incubate craving for sucrose, but not alcohol.

To align with our previous findings that female rats that displayed more active than passive coping strategies self-administered more alcohol after TMT exposure, we only included females in experiment 2 in the present study [18]. Further experiments will be needed to determine if the correlation between reduced lever responses for sucrose and active stress-coping behavior, uncovered here in females, is also present in males.

### 4.3 Conclusions and implications

The individual differences in stress-coping behavior, namely digging and immobility, displayed during TMT exposure in the present study align with our previous findings [9, 13, 18]. We expand on our previous study showing that more digging than immobility during predator odor exposure is associated with increased alcohol self-administration by showing digging is correlated with increased impulsive choice and reduced sucrose self-administration when sucrose and alcohol are both available. Additionally, we show that behavioral differences in impulsive choice and sucrose self-administration are present even prior to stressor exposure. In summary, more active stress-coping behavior like digging predicts susceptibility to stress-induced changes in reward-related decision making.

Interestingly, lever responses and motivation for sucrose were lower in high-digging rats in Experiment 2, but high-digging rats in Experiment 1 did not show any reduced behavior when learning and performing the delay discounting task for sucrose. Delay-discounting training involved a limited number of trials in all training phases, which may have occluded differences in motivation for sucrose because animals were food restricted, and almost all of them completed the maximum number of limited trials by the end of training. Results from Experiments 1 and 2 suggest that active stress coping (digging) is related to both pre-existing anhedonia-like behavior as well as susceptibility to stress-induced increases in impulsive choice.

Although anhedonia is more common in individuals with PTSD, it is unclear whether increased anhedonia occurs as a result of trauma or, alternatively, if anhedonia is a risk factor for development of PTSD [68, 69, 70]. Our results support the latter hypothesis and suggest that anhedonia may be a stable trait in a subset of individuals that display more active coping and are more susceptible to the negative effects of stress. Given that animal studies allow for the establishment of baseline anhedonia-like behavior before stressor exposure, future animal studies may be crucial to determine the relationship between anhedonia and susceptibility to stressors and the underlying neurobiology of this relationship.

We showed that high-digging rats displayed both more impulsive choice (Experiment 1) and reduced motivation for sucrose (Experiment 2), an indicator of anhedonia. Recent studies have shown associations between higher anhedonia-like behavior and steeper delay discounting in individuals with PTSD, but not in healthy controls [50, 71]. Therefore, high-digging rats may serve as a model of the subset of individuals more likely to develop PTSD after exposure to a major trauma/stressor, which adds validity to predator odor exposure as a model of PTSD. Human imaging suggests differences in reward-responsivity in the nucleus accumbens and functional connectivity between the nucleus accumbens and cortical regions are involved in the association between anhedonia-like behavior and impulsive choice [71]. Given the importance of the nucleus accumbens in reward processing and delay discounting [72], as well as evidence that projections from the medial prefrontal cortex to the nucleus accumbens mediate impulsive choice [73], differences in these brain regions and their connectivity could be evaluated in high-digging rats in future studies, particularly before stressor exposure.

Overall, these findings suggest that individual differences in decision-making behavior (impulsive choice for sucrose and motivation to self-administer sucrose) may predict the behavioral response to a stressful experience and, in turn, predict the susceptibility to negative effects of a stressor on future decision making. Future studies could examine if baseline measures of impulsive choice and anhedonia could be used to identify individuals at greater risk of developing psychopathology after a traumatic experience. These results also reveal complex relationships between decision making and behavioral responses to stress and provide a basis for future studies examining the overlapping neurobiological mechanisms of these relationships. Additionally, these findings highlight the importance of examining individual differences in stress responsivity by showing how effects in some individuals can be occluded by a lack of effect, or even an opposing effect, in other individuals.

## Acknowledgements

We would like to thank Dr. Christopher Lapish for graciously providing the delay discounting programs and code, and Dr. Lapish and Dr. Kathleen Bryant for analysis advice for delay discounting.

## Author Contributions (CRediT)

**BNB**: Conceptualization, methodology, formal analysis, investigation, data curation, writing - original draft, writing - review & editing, visualization, supervision, project administration. **MEH**: Investigation, formal analysis, data curation, writing - original draft, writing - review & editing. **CGK**: Formal analysis, data curation, writing - original draft, writing - review & editing. **HS**: Investigation, data curation. **JB**: Conceptualization, methodology, formal analysis, resources, writing - review & editing, visualization, supervision, project administration, funding acquisition.

