## Supplementary Material for "Stress-coping behavior during predator odor exposure is associated with differences in decision making"

### Supplementary Results

#### There were no effects of sex or digging group on behavior during training on delay discounting task

To ensure there were no differences in learning the delay discounting task between digging groups that could later influence delay discounting, we compared the averages of several behavioral measures throughout the phases of training between groups. For the average number of reinforcers retrieved of the maximum 30 delivered during magazine training, there were no main effects of digging group ($F_{(2, 32)}$=2.02, p = 0.149) or sex ($F_{(1, 32)}$=0.303, p = 0.586), and there was no interaction ($F_{(2, 32)}$=0.892, p = 0.420; 2-way ANOVA; Supplementary Fig S1A). For the average number of reinforcers earned of the maximum 100 during overnight training, there were no main effects of digging group ($F_{(2, 32)}$=0.603, p = 0.553) or sex ($F_{(1, 32)}$=0.0019, p = 0.966), and there was no interaction ($F_{(2, 32)}$=0.192, p = 0.827; 2-way ANOVA; Supplementary Fig S1B). For the average number of reinforcers retrieved during overnight training, there were no main effects of digging group ($F_{(2, 32)}$=2.282, p = 0.118) or sex ($F_{(1, 32)}$=0.113, p = 0.739), and there was no interaction ($F_{(2, 32)}$=0.493, p = 0.616; 2-way ANOVA; Supplementary Fig S1C). For the average number of reinforcers earned of the maximum 30 during trial training, there were no main effects of digging group ($F_{(2, 32)}$=1.924, p = 0.163) or sex ($F_{(1, 32)}$=0.028, p = 0.868), and there was no interaction ($F_{(2, 32)}$=0.006, p = 0.994; 2-way ANOVA; Supplementary Fig S1D). For the average number of reinforcers retrieved of the maximum 30 during trial training, there were no main effects of digging group ($F_{(2, 32)}$=1.874, p = 0.170) or sex ($F_{(1, 32)}$=0.035, p = 0.853), and there was no interaction ($F_{(2, 32)}$=0.0033, p = 0.997; 2-way ANOVA; Supplementary Fig S1E). For the average number of choice trials completed of the maximum 30 during 0-sec delay training, there were no main effects of digging group ($F_{(2, 32)}$=0.238, p = 0.790) or sex ($F_{(1, 32)}$=0.755, p = 0.391), and there was no interaction ($F_{(2, 32)}$=0.172, p = 0.843; 2-way ANOVA; Supplementary Fig S1F). For the average number of reinforcers earned during 0-sec delay training, there were no main effects of digging group ($F_{(2, 32)}$=0.956, p = 0.395) or sex ($F_{(1, 32)}$=0.973, p = 0.331), and there was no interaction ($F_{(2, 32)}$=1.508, p = 0.237; 2-way ANOVA; Supplementary Fig S1G). For the average number of reinforcers retrieved during 0-sec delay training, there were no main effects of digging group ($F_{(2, 32)}$=0.749, p = 0.481) or sex ($F_{(1, 32)}$=0.422, p = 0.521), and there was no interaction ($F_{(2, 32)}$=1.416, p = 0.258; 2-way ANOVA; Supplementary Fig S1H). For the average i value during the last 10 trials of the final 4 sessions of 0-sec delay training, there were no main effects of digging group ($F_{(2, 32)}$=1.186, p = 0.319) or sex ($F_{(1, 32)}$=2.054, p = 0.162), and there was no interaction ($F_{(2, 32)}$=1.046, p = 0.363; 2-way ANOVA; Supplementary Fig S1I). Overall, these results suggest that there were no pre-existing differences between males and females or between controls and future digging groups in learning the task.

**There were no effects of TMT exposure group or digging group on behavior during alcohol and sucrose self-administration training**

To establish that there were no differences between groups that later became controls or TMT-exposed rats, we evaluated single-sever training data and pre-TMT alcohol vs. sucrose choice self-administration. Single-lever alcohol and sucrose training are shown on the same graph but were analyzed separately. When comparing between controls and all TMT rats for single-lever alcohol training, there was a main effect of session on alcohol lever responses ($F_{(9, 198)}=8.610$, $p < 0.001$), but there was no effect of future TMT exposure group ($F_{(1, 22)}=1.913$, $p = 0.181$) or interaction ($F_{(9, 198)}=1.611$, $p = 0.114$; 2-way rmANOVA; Supplementary Fig S2A). For single-lever sucrose training, there was a main effect of session on sucrose lever responses ($F_{(2.465, 54.22)}=10.56$, $p < 0.001$), but there was no effect of future TMT exposure group ($F_{(1, 22)}=0.0003$, $p = 0.987$) or interaction ($F_{(2.465, 54.22)}=0.825$, $p = 0.466$; 2-way rmANOVA; Supplementary Fig S2A). When comparing between digging groups for single-lever alcohol training, there was a main effect of session on alcohol lever responses ($F_{(5.073, 106.5)}=10.44$, $p < 0.001$), but there was no effect of future digging group ($F_{(2, 21)}=0.919$, $p = 0.415$) or interaction ($F_{(10.15, 106.5)}=1.053$, $p = 0.405$; 2-way rmANOVA; Supplementary Fig S2B). For single-lever sucrose training, there was a main effect of session on sucrose lever responses ($F_{(2.574, 54.05)}=11.06$, $p < 0.001$), but there was no effect of future digging group ($F_{(2, 21)}=1.834$, $p = 0.184$) or interaction ($F_{(5.148, 54.05)}=1.510$, $p = 0.201$; 2-way rmANOVA; Supplementary Fig S2B).

After single-lever training, rats were trained in choice sessions to press one lever for alcohol and the other for sucrose. Alcohol preference (the ratio of alcohol lever responses to sucrose lever responses), alcohol lever responses, and sucrose lever responses were measured to determine if there were any pre-existing differences in choice prior to TMT exposure. For controls compared to all TMT rats, there was a main effect of session on alcohol preference ($F_{(6.053, 133.2)}=5.487$, $p < 0.001$), but there was no effect of future TMT exposure group ($F_{(1, 22)}=0.286$, $p = 0.598$) or interaction ($F_{(6.053, 133.2)}=1.234$, $p = 0.293$; 2-way rmANOVA; Supplementary Fig S2C). Similarly, when TMT rats were separated by digging group, there was a main effect of session on alcohol preference ($F_{(5.991, 125.8)}=6.264$, $p < 0.001$), but there was no effect of future digging group ($F_{(2, 21)}=0.342$, $p = 0.715$) or interaction ($F_{(11.98, 125.8)}=1.642$, $p = 0.088$; 2-way rmANOVA; Supplementary Fig S2D). For controls compared to all TMT rats, there was a main effect of session on alcohol lever responses ($F_{(6.061, 133.3)}=2.426$, $p = 0.029$), but there was no effect of future TMT exposure group ($F_{(1, 22)}=0.803$, $p = 0.380$) or interaction ($F_{(6.061, 133.3)}=0.747$, $p = 0.615$; 2-way rmANOVA; Supplementary Fig S2E). Similarly, when TMT rats were separated by digging group, there was a main effect of session on alcohol lever responses ($F_{(5.885, 123.6)}=2.926$, $p = 0.011$), but there was no effect of future digging group ($F_{(2, 21)}=1.766$, $p = 0.196$) or interaction ($F_{(11.77,$

$_{123.6)}$=0.809, p = 0.639; 2-way rmANOVA; Supplementary Fig S2F). Finally, for controls compared to all TMT rats, there was a main effect of session on sucrose lever responses ($F_{(3.909, 86.00)}$=7.922, p < 0.001), but there was no effect of future TMT exposure group ($F_{(1, 22)}$=0.0002, p = 0.988) or interaction ($F_{(3.909, 86.00)}$=1.196, p = 0.318; 2-way rmANOVA; Supplementary Fig S2G). Similarly, when TMT rats were separated by digging group, there was a main effect of session on sucrose lever responses ($F_{(3.656, 76.77)}$=9.255, p < 0.001 but there was no effect of future digging group ($F_{(2, 21)}$=0.0177, p = 0.983) or interaction ($F_{(7.311, 76.77)}$=1.866, p = 0.084; 2-way rmANOVA; Supplementary Fig S2H). These results suggest that although alcohol preference, alcohol lever responses, and sucrose lever responses changed across sessions as sucrose concentration was increased, there were no differences in pre-TMT choice learning based on future TMT exposure or behavior during TMT exposure.

### Supplementary Figures and Legends

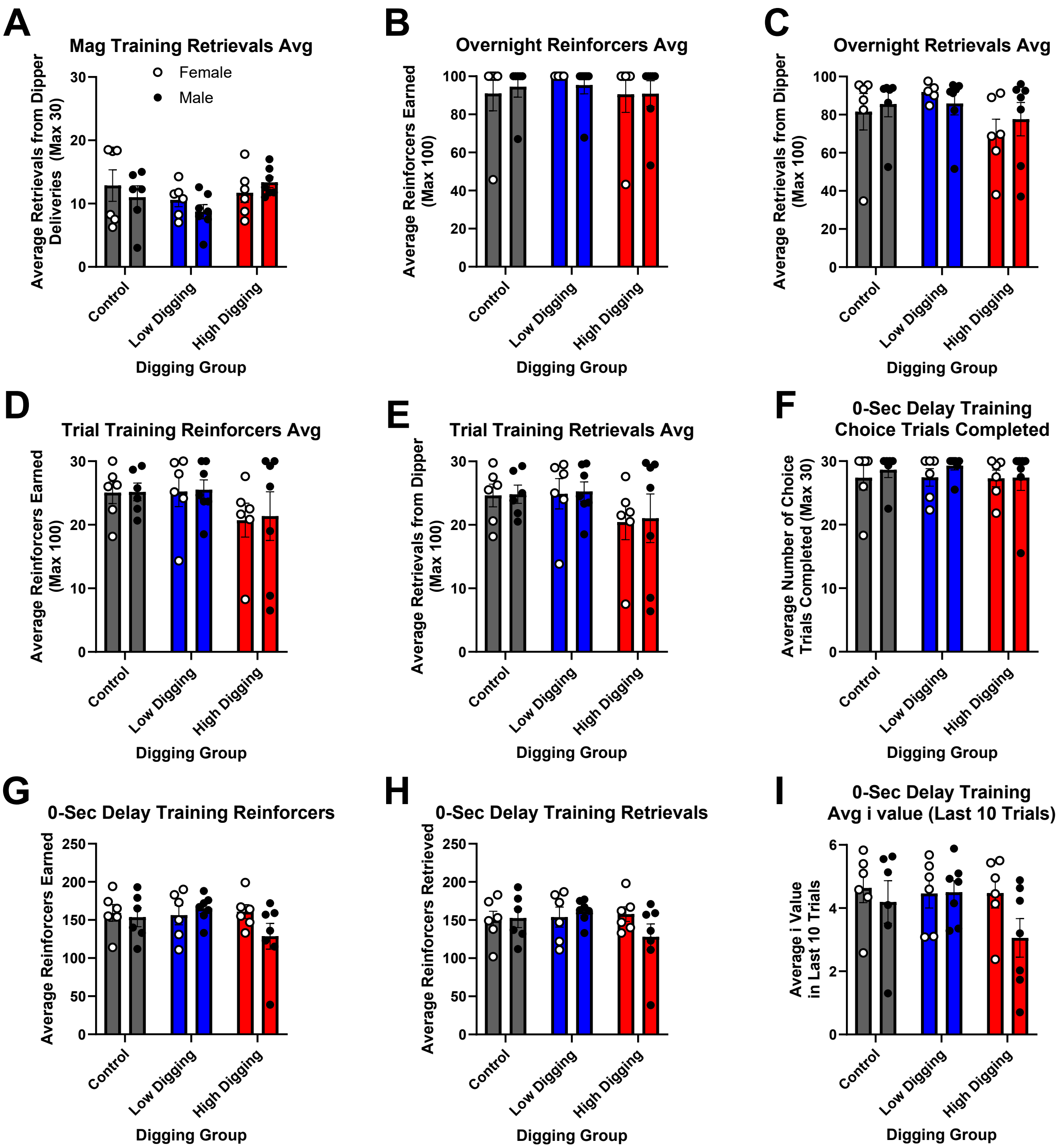


**Supplementary Figure 1**: There was no effect of sex or digging group on learning of delay discounting task prior to establishing delay discounting curve

The average of several measures during training phases were compared using sex and future TMT digging group as independent variables. There were no main effects or interactions for retrievals from dipper during magazine training (A); reinforcers earned (B) or retrieved (C) during overnight training; reinforcers earned (D) or retrieved (E) during trial training; choice trials completed (F), reinforcers earned (G) or retrieved (H), or average I value in the last 10 trials of the final 4 days (I) during 0-second delay training. Graphs show group means ± SEM, with females indicated by open symbols and males by close symbols.

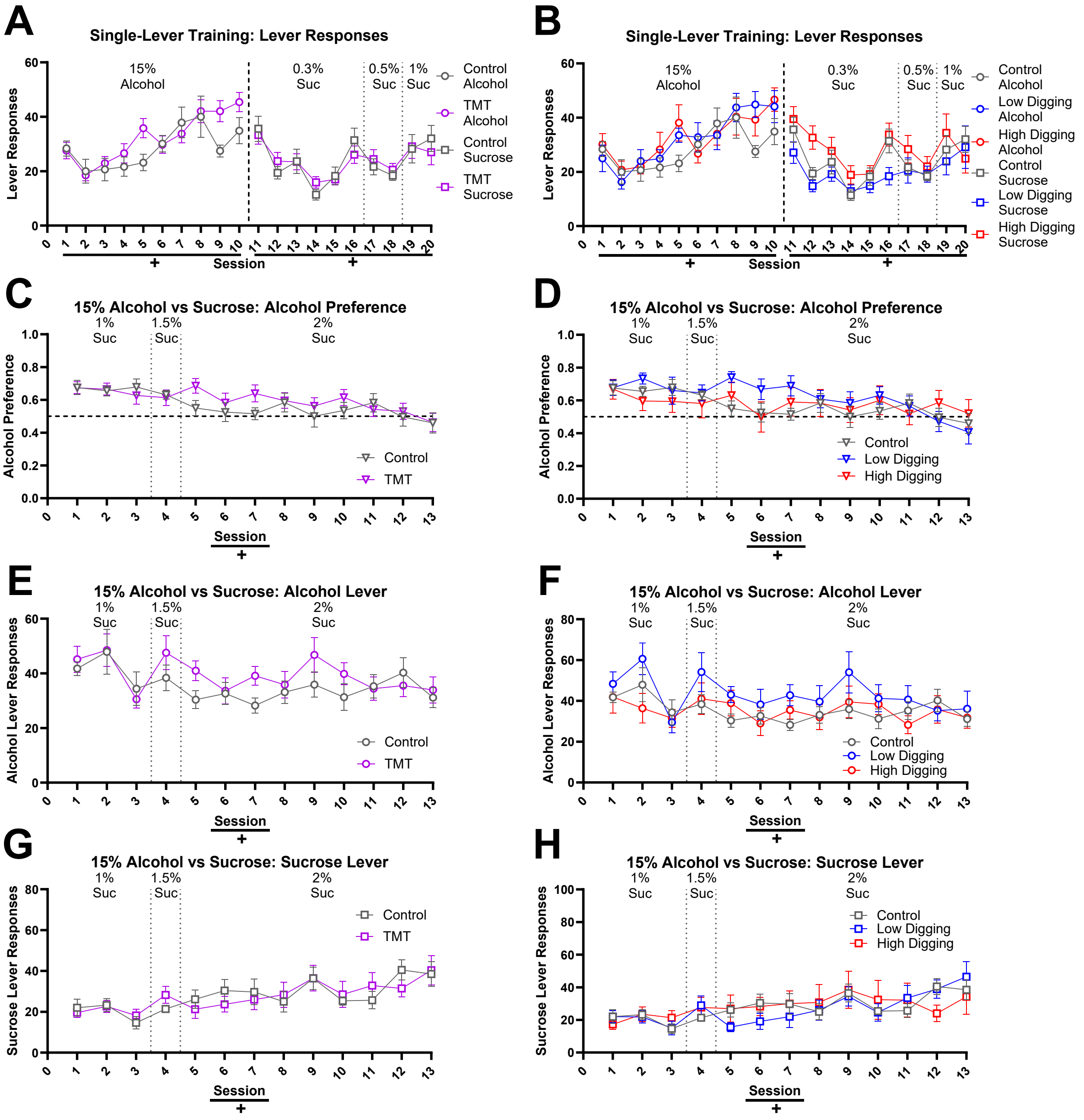


**Supplementary Figure 2**: There was no effect of TMT group or digging group on learning to self-administer alcohol and sucrose or on alcohol vs. sucrose choice prior to TMT exposure

After sucrose fading procedures for acquisition of alcohol self-administration, rats were trained first in single-lever sessions to self-administer 15% alcohol and then sucrose (0.3-

1%). Lever presses for alcohol and then sucrose in single-lever training sessions are shown for control vs. all TMT rats (A), and then between controls, low-digging, and high-digging rats (B). After single-lever training, rats underwent choice training, where rats could self-administer 15% alcohol using one lever and 1-2% sucrose using another lever. Alcohol preference (C-D), alcohol lever presses (E-F), and sucrose lever presses (G-H) are shown for controls vs. all future TMT-exposed rats (C, D, E) and between control and rats that became low-digging or high-digging rats during later TMT exposure (D, F, H). Main effects of session are indicated by +. Graphs show group means ± SEM.

### Supplementary Tables

| | | Pre-TMT | | | | | | | Post-TMT | | | | | |
|---|---|---|---|---|---|---|---|---|---|---|---|---|---|---|
| | Immobility | 0-sec i | 2-sec i | 4-sec i | 8-sec i | 16-sec i | AUC | k | 2-sec i | 4-sec i | 8-sec i | 16-sec i | AUC | k |
| **Digging** | r=-0.012 p=0.953 | r=-0.122 p=0.554 | r=-0.221 p=0.277 | r=-0.110 p=0.593 | r=-0.290 p=0.150 | r=-0.135 p=0.511 | r=-0.253 p=0.213 | r=0.260 p=0.200 | r=-0.184 p=0.367 | r=-0.527 **p=0.006*** | r=-0.436 **p=0.026*** | r=-0.004 p=0.985 | r=-0.469 **p=0.016*** | r=0.390 **p=0.049*** |
| **Immobility** | — | r=-0.050 p=0.809 | r=-0.010 p=0.960 | r=-0.195 p=0.339 | r=-0.290 p=0.354 | r=-0.049 p=-0.812 | r=-0.176 p=0.390 | r=0.142 p=0.489 | r=-0.332 p=0.098 | r=-0.058 p=0.778 | r=-0.212 p=0.299 | r=-0.213 p=0.296 | r=-0.260 p=0.199 | r=0.114 p=0.581 |

**Supplementary Table 1**: Correlations between stress-coping behavior and pre- and post-TMT measures of delay discounting

Significant correlations ($p<0.05$) are indicated by *; N=26.

| | | Pre-TMT | | | Post-TMT | | | | | | |
|---|---|---|---|---|---|---|---|---|---|---|---|
| | | Choice | | | Choice | | | PR Choice | | | |
| | Immobility | Alc Lev | Suc Lev | Alc Pref | Alc Lev | Suc Lev | Alc Pref | Alc Lev | Alc Breakpoint | Suc Lev | Suc Breakpoint |
| **Digging** | r=-0.612 **p=0.012*** | r=-0.193 p=0.473 | r=-0.502 **p=0.047*** | r=0.347 p=0.188 | r=-0.179 p=0.507 | r=-0.572 **p=0.021*** | r=0.310 p=0.243 | r=-0.148 p=0.585 | r=-0.189 p=0.483 | r=-0.440 p=0.088 | r=-0.526 **p=0.037*** |
| **Immobility** | — | r=0.183 p=0.497 | r=0.138 p=0.611 | r=-0.077 p=0.776 | r=-0.095 p=0.728 | r=0.549 **p=0.027*** | r=-0.408 p=0.116 | r=-0.113 p=0.678 | r=-0.069 p=0.798 | r=0.118 p=0.663 | r=0.231 p=0.389 |

**Supplementary Table 2**: Correlations between stress-coping behavior and pre- and post-TMT measures of alcohol and sucrose self-administration, preference, and motivation

Significant correlations ($p<0.05$) are indicated by *; N=16.
